# Stronger together: harnessing natural algal communities as potential probiotics for inhibition of aquaculture pathogens

**DOI:** 10.1101/2024.08.27.609935

**Authors:** Dóra Smahajcsik, Line Roager, Mikael Lenz Strube, Sheng-Da Zhang, Lone Gram

**Affiliations:** Department of Biotechnology and Biomedicine, Technical University of Denmark, DK-2800 Kgs. Lyngby, Denmark

**Keywords:** algal microbiome, probiotics, *Vibrio*, *Isochrysis galbana*, *Tetraselmis suecica*, aquaculture, pathogen antagonism

## Abstract

Intensive fish rearing in aquaculture is challenged by infectious diseases, and although vaccines have been successfully developed for mature fish, alternative disease control measures are needed for fish larvae and juveniles. Probiotics offer a promising alternative to antibiotics, with the potential to reduce the risk of antibiotic resistance. Probiotics are typically isolated and used as pure cultures, however, in natural environments it is the concerted effort of the complex microbiome that keeps pathogens at bay. Here, we developed an *in vitro* assay to evaluate the anti-pathogen efficacy of mixed algal microbiomes from the live-feed microalgae *Tetraselmis suecica* and *Isochrysis galbana.* The inhibition of a GFP-tagged *Vibrio anguillarum*, a key fish pathogen, by microbial communities, was measured and quantified as reduction in fluorescence. The *Isochrysis galbana* microbiome was more inhibitory to *V. anguillarum* than the *Tetraselmis suecica* microbiome. During co-culture with the pathogen, the bacterial density of the *Isochrysis* microbiomes increased whilst the diversity was reduced as determined by metataxonomic analyses. Bacteria isolated from the fully inhibitory microbiomes were members of *Alteromonadaceae, Halomonadaceae, Rhodobacteraceae, Vibrionaceae, Flavobacteriaceae,* and *Erythrobacteraceae*. Although some strains individually inhibited the pathogen, these were not the key members of the microbiome and enhanced inhibition was observed when *Sulfitobacter pontiacus* D3 and *Halomonas campaniensis* D2 were co-cultured, even though neither were inhibitory as monocultures. Thus, this study demonstrates that microbial communities derived from natural algal microbiomes can have anti-pathogen effects, and that bacterial co-cultures may offer synergistic advantages over monocultures as probiotics, highlighting their promise for aquaculture health strategies.

**IMPORTANCE:** Aquaculture is the fastest growing food protein producing sector and sustainable disease control measures are required. Probiotics have gained interest as a promising solution for combating fish pathogens and using mixtures of microorganisms rather than pure cultures may represent a more stable pathogen control. We developed an assay using GFP-tagging of a fish pathogen, enabling the quantitative assessment of the anti-pathogen effects of complex microbiomes. We show that the efficiency of pathogen suppression can be increased with co-cultures compared to monocultures, thus emphasising the potential in using mixtures of bacteria as probiotics.

## INTRODUCTION

As the human population continues to rise, sustainable production of dietary protein is becoming increasingly important. Aquaculture is a key sector in supplying such high quality protein and has for decades been the fastest growing protein producing food sector (1). However, intensive rearing of fish carries with it the risk of disease outbreaks, which are especially damaging during larval development (1, 2) where disease outbreaks can result in substantial economic losses as seen in a 1988 incident where vibriosis led to a 40% loss of juvenile turbot in a Norwegian hatchery (3, 4). Vibrionaceae infections pose a considerable challenge to marine larviculture and can be caused by a number of *Vibrio* species, including *V. anguillarum* that can infect up to 80 different fish species (5, 6). Despite advances in vaccine development, vibriosis remains a significant problem in aquaculture, especially due to fish larvae lacking a mature immune system, rendering vaccination ineffective (1, 2).

Many commercially valuable marine fish species require live feed, such as microalgae, at the larval stages since suitable artificial feed formulations are not available (7, 8), however, live feed can serve as the infection vector for pathogenic bacteria in larval rearing, leading to swift and widespread infections and complicating disease management (8, 9). Historically, antibiotics have been used to control these infections among larvae and juveniles, but since this may lead to spread of antimicrobial resistance (AMR) of both clinical and agri- and aquacultural relevance, alternative approaches are urgently needed (10). A promising strategy to mitigate bacterial pathogen proliferation in live feed and fish larvae is the use of probiotic bacteria which can inhibit pathogenic bacteria (11–13). Probiotic microorganisms, as defined by FAO and WHO, are “live microorganisms that, when administered in adequate amounts, confer a health benefit on the host” (14). As reviewed in Sonnenschein *et al.* (15), probiotics can be an environmentally sustainable and economically viable approach to counteract economic losses due to bacterial pathogens in aquaculture, and have been intensively studied in the past decades (2, 16). However, the vast majority of such studies have focused on utilising pure cultures of single probionts (17) which may pose challenges in terms of prolonged establishment of a probiont in live feed microbiomes (18). In contrast to traditional single-isolate probiotics, complex microbiomes may offer a more holistic approach by capitalising on the synergistic interactions among multiple microorganisms, including bacteria, to effectively combat pathogens (19). Several studies have focused on use of probiotics in larviculture and have suggested that deriving potential probiotic cultures from the aquaculture system offers an ecological advantage when they are re-introduced (11, 15, 20, 21).

Microalgae, such as *Tetraselmis suecica* and *Isochrysis galbana*, that are used as live feed in many larviculture productions are colonized by a microbiome that can influence the growth and metabolism of the algae (22–24). The algal microbiome depends on the host species (25), is taxonomically diverse and harbours a large bioactive potential (26, 27). The microbial world is characterized by an intricate web of interactions some of which are mediated by small molecules through which microorganisms exchange information with their neighbours, competitors, and hosts. These interactions shape the microbial community and its functionality, however, monocultures lack these interactions that are important for e.g. inducing expression of biosynthetic gene clusters producing bioactive compounds (28–30). We therefore hypothesize that the collective microalgal microbiome and not only pure cultures of probionts could serve to counteract invasion of aquaculture pathogens. Using a complex microbiome to inhibit a target bacterium, *Vibrio anguillarum*, we hope to harvest the enhanced inhibitory properties promoted by the microbiome interactions.

Whilst the study of single culture inhibition is relatively simple, the high throughput screening of pathogen inhibition by a complex microbiome is more challenging, and a purpose of the present study was to develop a simple screening assay that would allow determining pathogen inhibition by a complex microbiome. Subsequently we characterized the inhibitory microbiomes by metataxonomic analyses and tested mixtures of bacteria derived from these inhibitory microbiomes for their capability to inhibit the fish pathogen.

## MATERIALS AND METHODS

### Bacterial and algal strains and culture conditions

The fish pathogen *Vibrio anguillarum* strain NB10 with a fluorescent tag integrated into the large chromosome (NB10_pNQFlaC4-gfp27) (31) (later referred to as *Vibrio anguillarum* NB10_gfp) was used as target bacterium in the inhibition assays. The expression of GFPmut3* gene was controlled by a constitutive promoter P_A1/04/03_ (32). *Phaeobacter piscinae* strain S26 (21) was used as a known antagonist of *Vibrio* as a positive control for establishing the fluorescence-based screening assay. *P. piscinae* and *V. anguillarum* were stored at –80°C and routinely cultivated on Marine Agar (MA; Difco2216 BD) at 25°C or in Marine Broth (MB; Difco2216 BD) at 25°C and 200 rpm. MA supplemented with 4 μg/mL chloramphenicol was used to maintain *V. anguillarum* NB10_gfp. All bacterial cultures were stored and cultivated similarly, including *V. anguillarum* NB10 wild type (WT), and bacterial isolates obtained from algal cultures and enriched inhibitory microbiomes (see below).

Xenic microalgal cultures of *Tetraselmis suecica* and *Isochrysis galbana* were provided by aquaculture industry partners. Axenic algae *Isochrysis galbana* CCMP 1323 and *Tetraselmis suecica* CCAP 66/4 were purchased from the Bigelow National Center for Marine Algae and Microbiota (NCMA) and the Culture Collection of Algae and Protozoa (CCAP), respectively, and their axenic status determined by plating on MA. The algae were cultured in f/2 medium without silicate (f/2 – Si; NCMA; (33, 34) made with 3% Instant Ocean (IO; Aquarium Systems Inc., Sarrebourg, France) and cultivated at 18°C and constant illumination at ∼50 μE m^−2^ s^−1^. Cultures were maintained by continuous subculturing every 4 weeks, transferring 1% of the culture to 25 mL fresh medium.

### Fluorescence-based screening assay for GFP-labelled target organism

*V. anguillarum* NB10_gfp was grown overnight as described above and a 10-fold serial dilution was prepared using MB as a diluent. 180 μL of the dilutions were distributed to the wells of a sterile, black 96-well microplate with flat, clear bottom (Perkin Elmer, 6005430) in technical triplicates. *V. anguillarum* NB10 WT was included as a negative control for the fluorescence measurement, and sterile medium as a no-growth negative control. Fluorescence and absorbance were measured on a BioTek Cytation^TM^ 5 multi-mode microplate reader at 25°C with shaking (linear, continuous, 567 cpm). Measurements were taken every 20 minutes over 40 hours, measuring fluorescence (excitation: 485/15 nm, emission: 513/15 nm) and OD_600_.

### Fluorescence-based inhibition assay by *Phaeobacter piscinae* S26

*Phaeobacter piscinae* S26 was co-cultured with *V. anguillarum* NB10_gfp in the fluorescence-based inhibition assay. Overnight cultures of both bacteria were prepared, 10-fold serially diluted in sterile MB and 90 μL of each dilution of both bacterial cultures was loaded into sterile, black 96-well microplates allowing crossed combinations of the two strains at the different dilutions (total well-content of 180 μL). Dilutions of *P. piscinae* S26 were included as a negative control for the fluorescence measurement, and sterile medium as a no-growth negative control. Dilutions of *V. anguillarum* NB10_gfp were included as a positive control for fluorescence. Absorbance and fluorescence of the plates were measured as described above. Bacterial counts of the precultures were determined by 10-fold serial dilution and plating on MA. After the assay, wells showing no fluorescent signal (the inhibitory microbiomes) were 10-fold serial diluted and plated on MA supplemented with 4 μg/mL chloramphenicol to confirm decreased *Vibrio* counts.

### Inhibition of *Vibrio anguillarum* by complex algal microbiomes

Microbiomes from two xenic microalgae, *Tetraselmis suecica* and *Isochrysis galbana,* were co-cultured with *V. anguillarum* NB10_gfp using the fluorescence-based screening assay (Figure 1). Samples from axenic algal cultures were included as controls without microbiomes. Five algal cultures were tested in the assay described above. This included the two axenic cultures (AXT: axenic *Tetraselmis*; AXI: axenic *Isochrysis*), two xenic cultures of *Tetraselmis suecica* (NT) and *Isochrysis galbana* (NI) that had been re-cultivated in the lab since 2019 and 2020, respectively, and a newly acquired (2023) *Isochrysis galbana* culture (NNI) from an aquaculture collaborator. Cultures (200 mL) of the two microalgae, both axenic and xenic, were prepared by inoculating 10% stock culture to fresh media and incubating them at 18°C and constant illumination for five days reaching approx. 6 log algal cells mL^-1^. The *V. anguillarum* NB10_gfp strain was grown overnight, and algal and bacterial counts were determined at the beginning of the experiment. Bacterial counts were determined by 10-fold serial dilution and plating on MA. Plates were incubated for two days for *V. anguillarum* and for four days for the bacterial counts from algal cultures. All CFU counts were log-transformed before further data handling and plotting. Algal cell counts were determined using microscopy and a Neubauer-improved chamber. To determine if any pathogen inhibition was caused by the algae or by their microbiomes, the xenic algal cultures were divided into two 100 mL fractions: the full culture (FC) with algal and bacterial cells, and a filtered microbiome (FM) fraction, where algal cells were removed by filtering (47-mm Ø, pore size 3-µm, mixed cellulose ester, Advantec A300A047A). Before filtering, the cultures were vortexed vigorously to detach bacteria from the algal cells.

**Figure 1.**
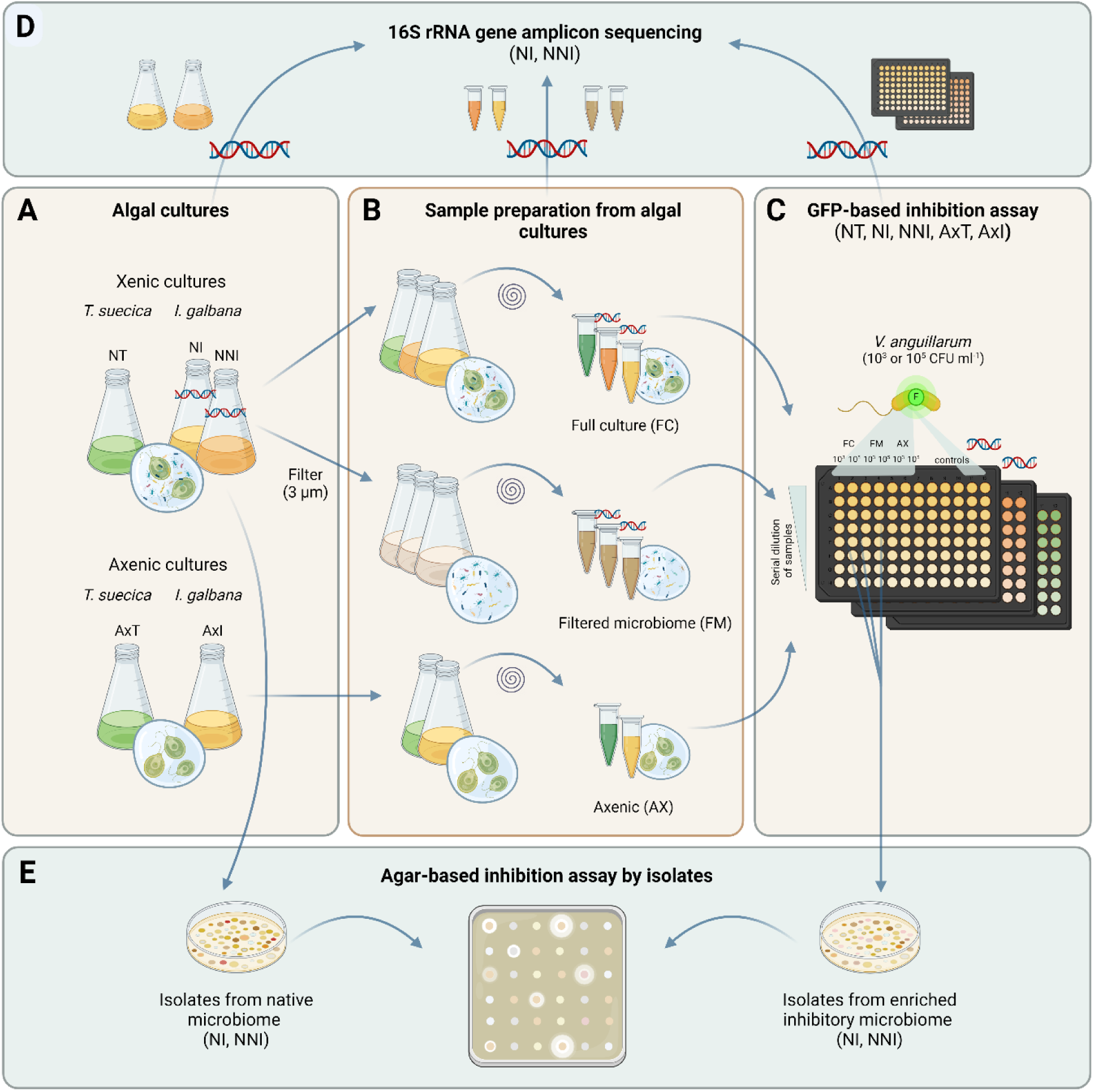
Yellow panels: Overview of algal cultures used in the *Vibrio* inhibition assays. Xenic cultures: *Tetraselmis suecica* (NT) and *Isochrysis galbana* (NI – continuously subcultured, NNI – freshly recruited microbiome). Axenic cultures: *T. suecica* (AxT) and *I. galbana* (AxI) with no microbiome (A). Sample preparation: Xenic cultures were divided into Full culture (FC, containing algal and bacterial cells) and Filtered microbiome (FM, algal cells removed by filtration) (B). All samples were 100-fold up-concentrated, then 10-fold serial dilutions were co-cultured with *Vibrio anguillarum* NB10_gfp (starting inoculum 10^3^ or 10^5^ CFU mL^-1^) in a GFP-based inhibition assay (C). Teal panels: NI and NNI samples indicated by DNA symbol were analysed through 16S rRNA gene amplicon sequencing (D). Bacterial isolates from the native microbiome and the enriched inhibitory microbiome of *I. galbana* (NI, NNI) were tested for inhibition of *V. anguillarum* in an agar-based assay (E). (Created by BioRender).

To confirm that marine viruses or viral particles did not influence the observed inhibition, the possible pathogen inhibition was tested with a filtered microbiome (FM) and a cell-free supernatant (cf-SN) of NT and NI samples. To prepare the cell-free supernatant, a 100 mL fraction of the FM samples was filtered through 0.22 µm polyethersulfonate (PES) filters (Nalgene™ Rapid-Flow™).

To increase the bacterial density of the microbiome (target ∼ 7 – 8 log CFU mL^-1^), 100 ml of algal cultures were harvested (2,445 x g, 10 min, 20°C) and resuspended in 1 mL sterile 3% IO. 10-fold serial dilutions of both up-concentrated algal fractions (FC and FM; 3% IO as diluent), the cell-free supernatants (cf-SN; 3% IO as diluent) and *Vibrio* cultures (MB as diluent) were prepared, and loaded into sterile, black 96-well microplates. Combinations of dilutions of the pathogen (target: 3 log CFU mL^-1^ and 5 log CFU mL^-1^) and algal samples (10^-1^ to 10^-5^ dilutions) were loaded. Algal cultures with no addition of the GFP tagged pathogen were included as negative control for the fluorescence measurement, and sterile medium as a no-growth negative control. Different dilutions of the pure culture of *V. anguillarum* NB10_gfp were included as a positive control. Fluorescence and OD_600_ were measured as described above. FM samples were measured on a different instrument (BioTek Synergy HTX) than all other samples, resulting in a different scale of the fluorescence measurements. To confirm pathogen inhibition after the assay, wells with fluorescence below 5% of the non-inhibited Vibrio control were 10-fold serial diluted and plated on chloramphenicol-supplemented plates (MA). The overview of sample preparation and experimental setup are summarized in Figure 1.

### Characterizing the enriched inhibitory microbiomes

During the co-culture inhibition assay, the bacterial density of the algal microbiomes was increased. Several dilutions of the algal microbiomes inhibited growth of *V. anguillarum* in a “dose dependent” manner and in all co-cultures, the bacterial density of the algal microbiomes increased. These microbiomes which caused suppression of the pathogen will be referred to as enriched inhibitory microbiomes. DNA was extracted from a selection of the wells taking samples from wells exhibiting a complete inhibition and wells allowing only marginal growth of *V. anguillarum*. Also, samples were taken from no-growth negative controls of the experiment, resulting in a total of 41 samples. Upon completion of the assay, the plates and samples were stored at −80°C until further processing, and then thawed at 4°C and samples kept on ice. 2 x 200 μL lysis buffer (400 mM NaCl, 750 mM sucrose, 20 mM EDTA, 50 mM Tris-HCl, 1 mg/mL lysozyme [pH 8.5]) was added to the wells. The contents of the wells were then resuspended and transferred to Eppendorf tubes and incubated at 37°C for 30 min. Proteinase K was added to a final concentration of 100 mg/mL and sodium dodecyl sulphate (SDS) to a final concentration of 1% vol/vol, and samples were then incubated overnight at 55°C, 300 rpm (35). The order of handling of samples was then randomized before extraction. DNA extraction was done by the Promega Maxwell 16 using the Maxwell LEV Blood DNA Kit, and the quality and purity of the DNA were measured using a DeNovix 439 DS-11+ spectrophotometer (DeNovix Inc., Wilmington, DE, USA) and confirmed by gel electrophoresis.

The V3-V4 region of the 16S rRNA gene of all samples was amplified by PCR as described in Roager *et al*. (2023) (36), using the primer set 341f (59-CCTACGGGNGGCWGCAG-39) and 805r (59-GACTACHVGGGTATCTAATCC-39) tagged with 30 unique octametric barcodes (Table S1). PCR products were purified using Agencourt AMPure XP magnetic beads (0.8:1 bead volume to DNA solution; Agencourt Bioscience Corporation, Beverly, MA, USA). Concentrations of the amplicons were determined using a Qubit 2.0 Fluorometer with the high sensitivity (HS) assay kit (Qubit^TM^ dsDNA HS assay; Invitrogen by Thermo Fisher Scientific Inc., Eugene, OR, USA). Samples were pooled in equimolar ratios before library preparation and sequencing on an Illumina NovaSeq 6000 platform with paired-end 250-bp reads (Novogene; Cambridge, United Kingdom).

### Analyses of sequencing data

Sequencing data analyses were performed using QIIME 2 version 2023.5 (37) and R version 4.2.1 (38). The raw reads were imported to QIIME2, demultiplexed, and primers and barcodes were removed using the cutadapt demux-paired plugin (39). Reads were then denoised and dereplicated using the DADA2 package (40) and an amplicon sequence variant (ASV) table was created leaving out sequences shorter than 400bp. A phylogenetic tree was constructed using MAFFT (41) and FastTree (42), via the phylogeny plugin. Alpha- and beta-diversity metrics were computed using the diversity plugin (43) at an even sampling depth of 26,226. A classifier trained on the Silva database 138 (44) and the specific primer sequences employed as mentioned above was used for taxonomic classification of the reads via the feature-classifier plugin (45, 46). ASVs which were either not assigned to a phylum, classified as chloroplasts/mitochondria or had a frequency < 10 were filtered out before further analysis in R. Two sterility controls were excluded from the dataset before further analysis. QIIME 2 artifacts were imported to R v4.2.1 using the qiime2R package (47) and analysis was continued in RStudio (48) with the tidyverse (49) package. For multivariate analyses using the vegan package, the ASV table was normalized by square-root transformation and Wisconsin standardization (50). PERMANOVA analysis was then performed using the beta-group-significance function of Qiime2 and the adonis plugin (51) in R with 999 permutations to test significant differences in beta diversity as a function of source culture (NI and NNI), degree of pathogen inhibition, microbiome dilution and sample fraction (FC and FM), based on Bray-Curtis dissimilarities calculated using the normalized ASV table. The analysis was conducted both on the complete dataset and on subsets divided by samples before and after the enrichment assay. Interactive effects were included in the models but were removed due to lack of significance.

### Isolation and identification of bacterial strains from *Isochrysis galbana* microbiomes

Culturable bacterial isolates were obtained from a subset of the enriched inhibitory *I. galbana* microbiomes (NI and NNI) analysed for their microbiome composition. Culture from wells where *V. anguillarum* was inhibited by the microbiomes was serially diluted and plated on MA. Isolates were picked representing colonies with different morphology. Also, strains were picked from the plating of the algal culture before testing inhibition (native microbiome), resulting in a total of 64 bacterial isolates. The strains were re-streaked for purity and glycerol stocks (20% v/v) were made and stored at −80°C. The isolates were later identified by 16S rRNA gene amplification by colony PCR using universal primers 27F and 1492R and subsequent Sanger sequencing by Macrogen (Amsterdam, Netherlands). Sequencing results were compared against the NCBI nucleotide database using the BLASTn function (52) to determine the putative bacterial taxa.

### Inhibition of *V. anguillarum* by pure cultures and co-cultures of bacterial isolates

Antagonism of pure cultures of the 64 bacterial isolates against *V. anguillarum* NB10_gfp was tested using an agar-based assay (53) using 50 μL of culture to 50 ml of agar and spotting 5 μL of an overnight culture of the isolates on the pathogen-embedded plates. Plates were incubated at 25°C and visually inspected for clearing zones after 24 and 48 hours. In a similar manner, dual cultures of isolates from the enriched inhibitory microbiome were spotted on embedded *Vibrio* and inspected for inhibition. Five isolates were chosen for investigating co-culture effects as representatives of the five non-*Vibrio* genera isolated from the enriched inhibitory *I. galbana* microbiomes (Table 2) with complete inhibition of 5.3 log CFU mL^-1^ starting inoculum of *V. anguillarum*. The strains were grown in liquid cultures (MB, 25°C, 200 rpm) for 24 hours and then mixed 1:1 in pairs. 10 μL of the mixtures were spotted on the pathogen-embedded plates. The plates were incubated at 25°C and visually inspected for clearing zones after 24 and 48 hours. Monocultures of the individual strains mixed 1:1 with MB were used as controls. To validate the results by the co-culture of isolates D2 and D3, their inhibition was tested against a selection of nine strains of *Vibrio anguillarum* with different virulence against fish larvae (54). Monocultures and co-cultures were spotted on the pathogen-embedded plates in biological triplicates, as described above. Biological triplicates of isolate H2 (*Phaeobacter piscinae*) were used as a positive control.

### Whole genome sequencing, annotation and analysis

The five isolates selected for the co-culture study were whole genome sequenced with Oxford Nanopore sequencing, using the GridION platform. Genomic DNA was extracted from overnight cultures with the NucleoSpin® Tissue Mini kit (Macherey-Nagel, 740952.5) protocol for bacteria, with three hours of incubation during pre-lysis. The quality, purity, and concentration of the DNA was assessed as described for amplicon sequencing. The library was prepared using the Rapid Barcoding Kit 24 V14 (SQK-RBK114.24). The resulting reads were processed by the dragonflye pipeline (55). Briefly, adaptors were trimmed using Porechop and reads smaller than 1000bp were removed with Nanoq. The genomes were then assembled with Flye and polished with Medaka. The assemblies were then annotated with Bakta v1.8.2 (56) and profiled for BGCs with antiSMASH 7.0 (57). Subsequently, the assembled genomes were taxonomically classified using autoMLST (58).

### Data availability

The sequencing data from the microbiome analysis and the assembled whole genomes are available on the Sequencing Read Archive under BioProject ID PRJNA1150956.

## RESULTS

### Fluorescence-based inhibition assay for GFP-labelled *Vibrio anguillarum*

*V. anguillarum* NB10_gfp, as expected, grew well in MB reaching an absorbance of 0.929 ± 0.003 at 600 nm (in 19 hours for 5.3 log CFU mL^-1^ starting inoculum). The starting inocula of *V. anguillarum* NB10_gfp ranged between 1.3 to 9.3 log CFU mL^-1^. In pure culture, the growth of *V. anguillarum* could be followed by the fluorescent signal indicating the same lag phase, growth rate and stationary phase entry as seen on the OD_600_ measurements (Figure 2A and 2B). *V. anguillarum* was then co-cultured with a dilution series of a known antagonist, *Phaeobacter piscinae* S26 in biological triplicates, to provide proof-of-concept that growth-inhibition of the pathogen could be detected by measuring (lack of) fluorescence signal of a fluorescently tagged target organism. The starting inocula of *P. piscinae* S26 ranged between 9.1 to 1.1 log CFU mL^-1^. As expected, lower starting concentrations of *Vibrio* were more easily inhibited than higher concentrations (data not shown) and, conversely, higher concentrations of *Phaeobacter* were more inhibitory than lower concentrations (Figure 2C). Quantifying *Vibrio* on chloramphenicol-supplemented plates after the assay, from the wells with fluorescent signal < 5% of the *Vibrio* positive control, confirmed inhibition of the pathogen (< 4,34 ± 0,63 log CFU/mL).

**Figure 2.**
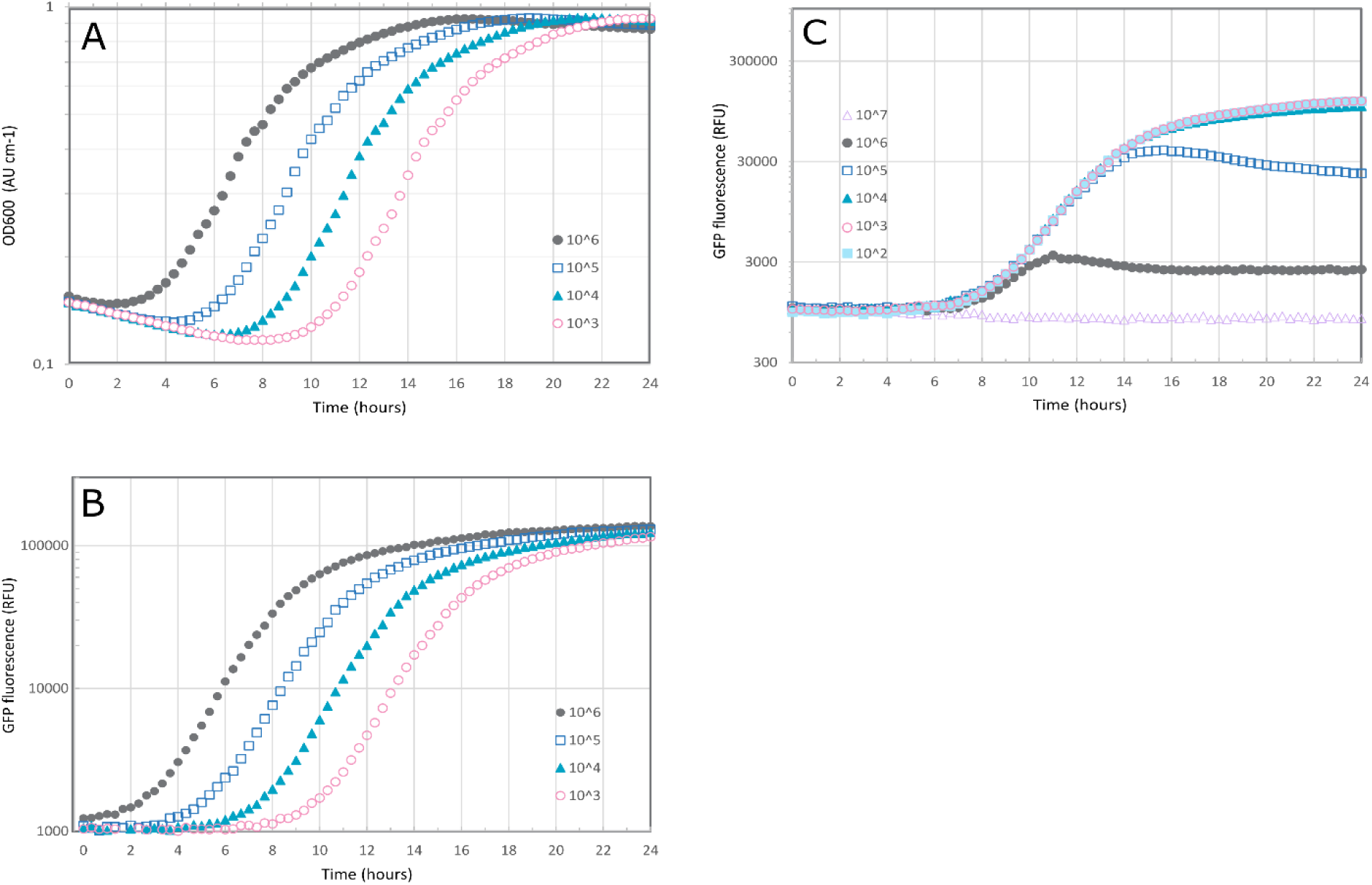
*Vibrio anguillarum* NB10_gfp at different initial concentrations: 10^6^ 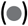, 10^5^ 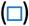, 10^4^ 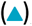, 10^3^ 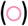 CFU mL^-1^, as measured by (A) absorbance at 600 nm and by (B) fluorescence (GFP) using excitation of 485/15 nm and emission of 513/15 nm, and the (C) inhibition profile of *V. anguillarum* NB10_gfp (initial concentration: 10^4^ CFU mL^-1^) by antagonist *Phaeobacter piscinae* S26 at different initial concentrations: 10^7^ 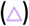, 10^6^ 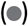, 10^5^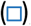, 10^4^ 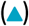, 10^3^ 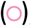, 10^2^ 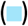 CFU mL^-1^, as measured by fluorescence (GFP, excitation: 485/15 nm, emission: 513/15 nm).

### Anti-pathogen inhibitory effect of algal microbiomes on *Vibrio anguillarum*

To investigate the inhibitory effect of their microbiomes on *V. anguillarum* NB10, dilutions of samples from cultures of two xenic microalgae, *I. galbana* and *T. suecica,* were co-cultured with the GFP-tagged fish pathogen. In the experiments, the starting inocula of *V. anguillarum* NB10_gfp was 3.1 ± 0.3 log CFU mL^-1^ (target: 3 log CFU mL^-1^) and 5.1 ± 0.3 log CFU mL^-1^ (target: 5 log CFU mL^-1^). Initial algal cell counts varied between 5.2 and 6.4 log cells mL^-1^ and up-concentration resulted in approx. 1.5 log increase to between 7.2 and 8.0 log cells mL^-1^. Initial bacterial counts in the *Isochrysis* samples varied between 5.7 and 6.6 log CFU mL^-1^ and in the up-concentrated FC fraction between 7.6 and 8.4 log CFU mL^-1^, whereas in the up-concentrated FM fraction between 7.6 and 8.1 log CFU mL^-1^ (Table S2). Microbiomes from both *T. suecica* and *I. galbana* inhibited growth of *V. anguillarum*, with the *I. galbana* microbiome being more potent (Figure 3A vs. 3D and 3B vs. 3E). The same inhibition pattern was observed for the filtered microbiomes (algal cells removed) as for the full culture (algal and bacterial cells) (Figure 3A vs. 3B and 3D vs. 3E). No inhibition was observed from samples from axenic algae cultures (Figure 3C and 3F) or cell-free supernatants (data not shown), indicating that it is indeed the microbiomes growing in the wells that inhibit the pathogen. Plating on chloramphenicol-supplemented plates from wells with decreased fluorescent signal confirmed that if the GFP signal was less than 10% of the control, the *Vibrio* was inhibited, with an abundance of < 5.67 ± 0.75 log CFU/mL, as opposed to approximately 9 log CFU/mL in the non-inhibited control. OD_600_ measurements can be seen in Figure S1.

**Figure 3.**
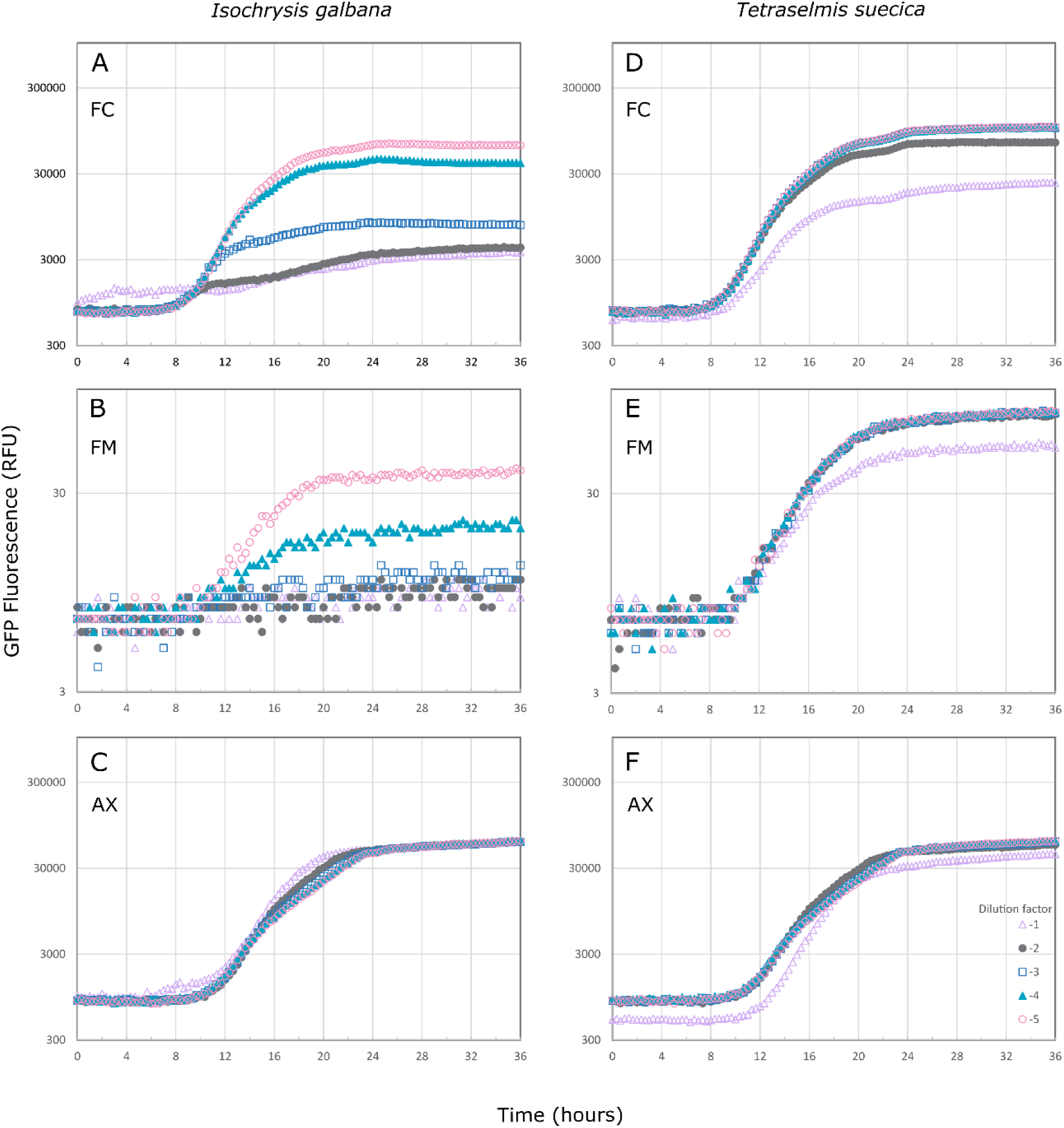
Inhibition by *Isochrysis galbana* (left) and *Tetraselmis suecica* (right) microbiomes of *Vibrio anguillarum* NB10_gfp (starting inoculum of 3.1 ± 0.3 log CFU mL^-1^), as measured by fluorescence (GFP, excitation: 485/15 nm, emission: 513/15 nm). The inhibitory effect of serial dilutions (10^-1^ 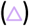, 10^-2^ 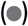, 10^-3^ 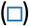, 10^-4^ 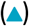, 10^-5^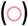) of different fractions of the algal cultures have been tested: full culture (FC; 3A and 3D), with algal and bacterial cells; filtered microbiome (FM; 3B and 3E), where algal cells have been removed; and axenic (AX; 3C and 3F) cultures, where the algae cells are free of bacteria. FM samples were measured on a different instrument, hence a different scale of RFU.

### Sequence based analysis of microbiomes

In total, 70,542,064 paired end reads were obtained from the 16S rRNA gene amplicons, with a mean of 1,500,894 reads per sample. After denoising, merging, filtering out chloroplasts and mitochondria, removal of short sequences (< 400 bp) and low abundant ASVs (total frequency < 10), this resulted in a total of 320 amplicon sequence variants (ASVs), with a mean read count per sample of 209,507 (Table S3).

### Microbiome composition and diversity analyses

The native microbiomes of *I. galbana* were dominated by *Alteromonadaceae, Rhizobiaceae, Flavobacteriaceae, Rhodobacteraceae* and *Hyphomonadaceae*. The microbiome composition of the two *I. galbana* cultures (NI and NNI) differed significantly (PERMANOVA, R^2^ = 0.312, p = 0.001), with NNI representing a relatively freshly recruited microbiome and NI representing a semi-adapted microbiome, subcultured many times in the laboratory over four years. The two cultures shared several genera, such as *Flavobacteriaceae, Rhizobiaceae, Alteromonadaceae* and *Rhodobacteraceae*, but also had distinct microbiome compositions. *Hyphomonadaceae* and *Cyclobacteriaceae* were unique to NI, and each culture had several distinct representatives of *Flavobacteriaceae* and *Rhodobacteraceae* (Figure S2).

The enriched inhibitory microbiomes sampled after the *Vibrio* inhibition assay were largely dominated by *Alteromonadaceae* and *Rhodobacteraceae*, with a higher relative abundance of *Vibrionaceae* compared to the native microbiome. As expected, the relative abundance of *Vibrionaceae* was highest in samples with marginal growth of the pathogen, compared to those with complete inhibition of the target pathogenic vibrio. However, *Vibrionaceae* were also detected in lower relative abundances in enriched cultures that fully inhibited *V. anguillarum*, suggesting these sequences could represent growth of native vibrios of the algal microbiome. The relative abundances of *Alteromonadaceae* and *Rhodobacteraceae* increased during the inhibition assay, whereas those of *Rhizobiaceae*, *Flavobacteriaceae*, and *Hyphomonadaceae* decreased.

Community richness (Faith index), diversity (Shannon index), and evenness (Pielou’s index) showed no significant differences between microbiomes with different degrees of pathogen inhibition (p > 0.05). However, based on a pairwise Kruskal-Wallis analysis, significant differences in Pielou’s index were observed between the NI and NNI cultures (p = 0.0041), with the NI culture having a higher diversity and evenness. PERMANOVA analysis of the full dataset indicated a significant shift in microbial community composition from before (native microbiome) to after the experiment (enriched inhibitory microbiome). Therefore, the data was split into two subsets for further analysis.

Post-enrichment, community composition was significantly affected by the source culture (NI vs. NNI) (PERMANOVA, R^2^ = 0.253, p = 0.001), degree of pathogen inhibition (PERMANOVA, R^2^ = 0.064, p = 0.017), microbiome dilution (PERMANOVA, R^2^ = 0.098, p = 0.001), and algal fraction (full culture (FC) vs. filtered microbiome (FM)) (PERMANOVA, R^2^ = 0.060, p = 0.025). Residual variation accounted for 52.5% of the total variation.

In conclusion, the degree of pathogen inhibition was associated with differences in the taxonomic composition of the sampled *Isochrysis* microbiomes. However, most variation between microbiomes was explained by differences between the two source cultures (NI and NNI).

### Isolation and identification of bacterial strains from *Isochrysis galbana* microbiomes

A total of 64 isolates were obtained from *I. galbana* microbiomes either from the native algal culture (19 strains) or from the enriched inhibitory microbiomes after *Vibrio* inhibition (45 strains). These were tentatively identified using full-length 16S rRNA gene sequencing (Table S4). Strains from the native microbiome were identified as members of the *Alteromonas* (2 isolates)*, Croceibacter* (1 isolate)*, Mameliella* (1 isolate)*, Phaeobacter* (3 isolates)*, Qipengyuania* (3 isolates)*, Roseovarius* (2 isolates)*, Ruegeria* (1 isolate) and *Sulfitobacter* (6 isolates) genera (Table 1), whereas strains from the enriched inhibitory microbiomes were identified as members of the *Alteromonas* (5 isolates)*, Halomonas* (1 isolate)*, Phaeobacter* (18 isolates)*, Roseovarius* (7 isolates)*, Sulfitobacter* (11 isolates) and *Vibrio* (3 isolates) genera (Table 2).

**Table 1.** Identification (16S rRNA gene sequence and BLAST) of bacterial isolates from the native *Isochrysis galbana* cultures.

| Top BLAST hit (NCBI) | Genus | Family | Number of<br>isolates | Present in<br>enrichment culture |
| --- | --- | --- | --- | --- |
| *Alteromonas* sp. | *Alteromonas* | *Alteromonadaceae* | 2 | Yes |
| *Croceibacter* sp. | *Croceibacter* | *Flavobacteriaceae* | 1 | No |
| *Mameliella* sp. | *Mameliella* | *Rhodobacteraceae* | 1 | No |
| *Phaeobacter* sp. | *Phaeobacter* | *Rhodobacteraceae* | 3 | Yes |
| *Qipengyuania* sp. | *Qipengyuania* | *Erythrobacteraceae* | 3 | No |
| *Roseovarius nubinihibens* | *Roseovarius* | *Rhodobacteraceae* | 2 | Yes |
| *Ruegeria* sp. | *Ruegeria* | *Rhodobacteraceae* | 1 | No |
| *Sulfitobacter* sp. | *Sulfitobacter* | *Rhodobacteraceae* | 6 | Yes |

**Table 2.**
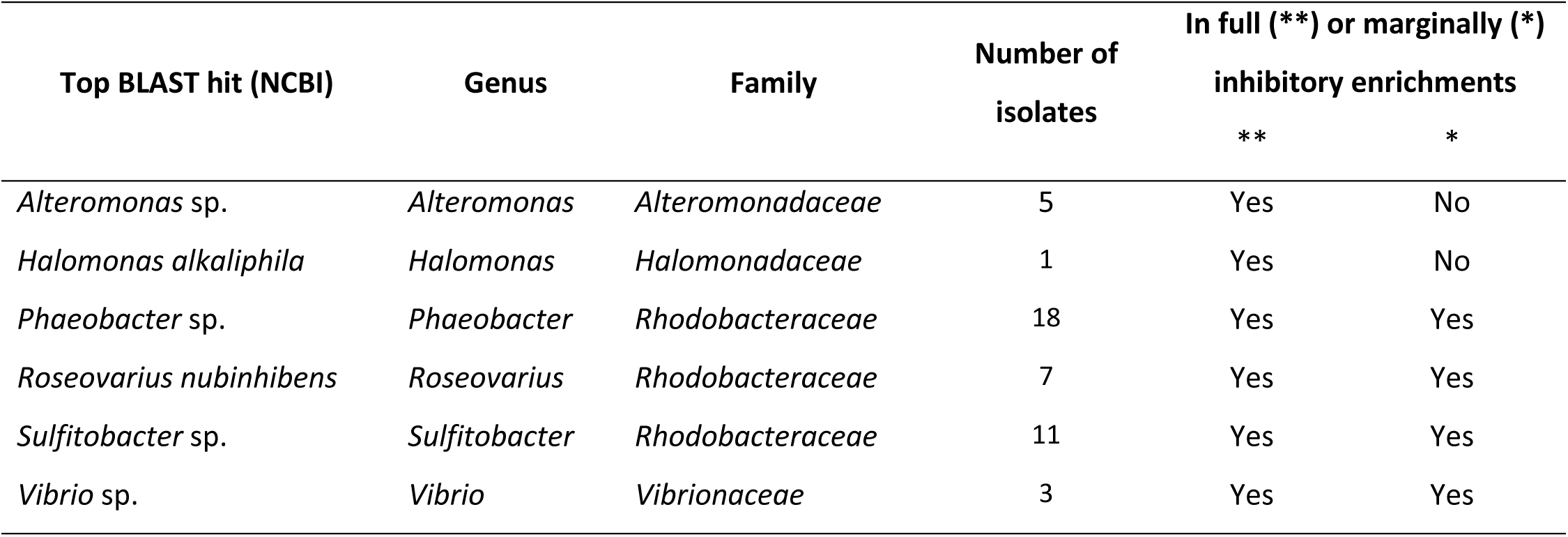
Identification (16S rRNA gene sequence and BLAST) of bacterial isolates from the enriched inhibitory *Isochrysis galbana* cultures after co-culture with *Vibrio anguillarum* NB10_gfp.

### Inhibition of *Vibrio anguillarum* by pure cultures and co-cultures of bacterial isolates

Pure cultures of all 64 bacterial isolates were tested for their ability to inhibit *V. anguillarum* NB10 and 17 isolates with pronounced inhibitory activity (zone of inhibition in the embedded pathogen) were all identified as *Phaeobacter* sp. (Table 3, Figure S3). The dual co-cultures of representative isolates showed inhibition zones in all co-cultures that included *Phaeobacter piscinae* (isolate H2). In some cases, the co-culture of *P. piscinae* (isolate H2) and *Roseovarius* sp. (isolate C7) resulted in a larger clearing zone as compared to the monoculture of the *P. piscinae*, however this result was not consistently reproducible. Of special interest was the co-culture of *Sulfitobacter pontiacus* (isolate D3) and *Halomonas campaniensis* (isolate D2), which despite showing only slight (*Halomonas campaniensis)* or no (*Sulfitobacter pontiacus)* inhibition in single culture, were remarkably inhibitory when grown together (Figure 4). Monocultures and co-cultures of these two isolates were subsequently tested against eight additional *V. anguillarum* strains with different virulence and the co-culture showed varying degrees of inhibition against *V. anguillarum* 775, 4299, and PF7 (Table S5). Isolate H2 showed strong inhibition against all *V. anguillarum* strains.

**Figure 4.**
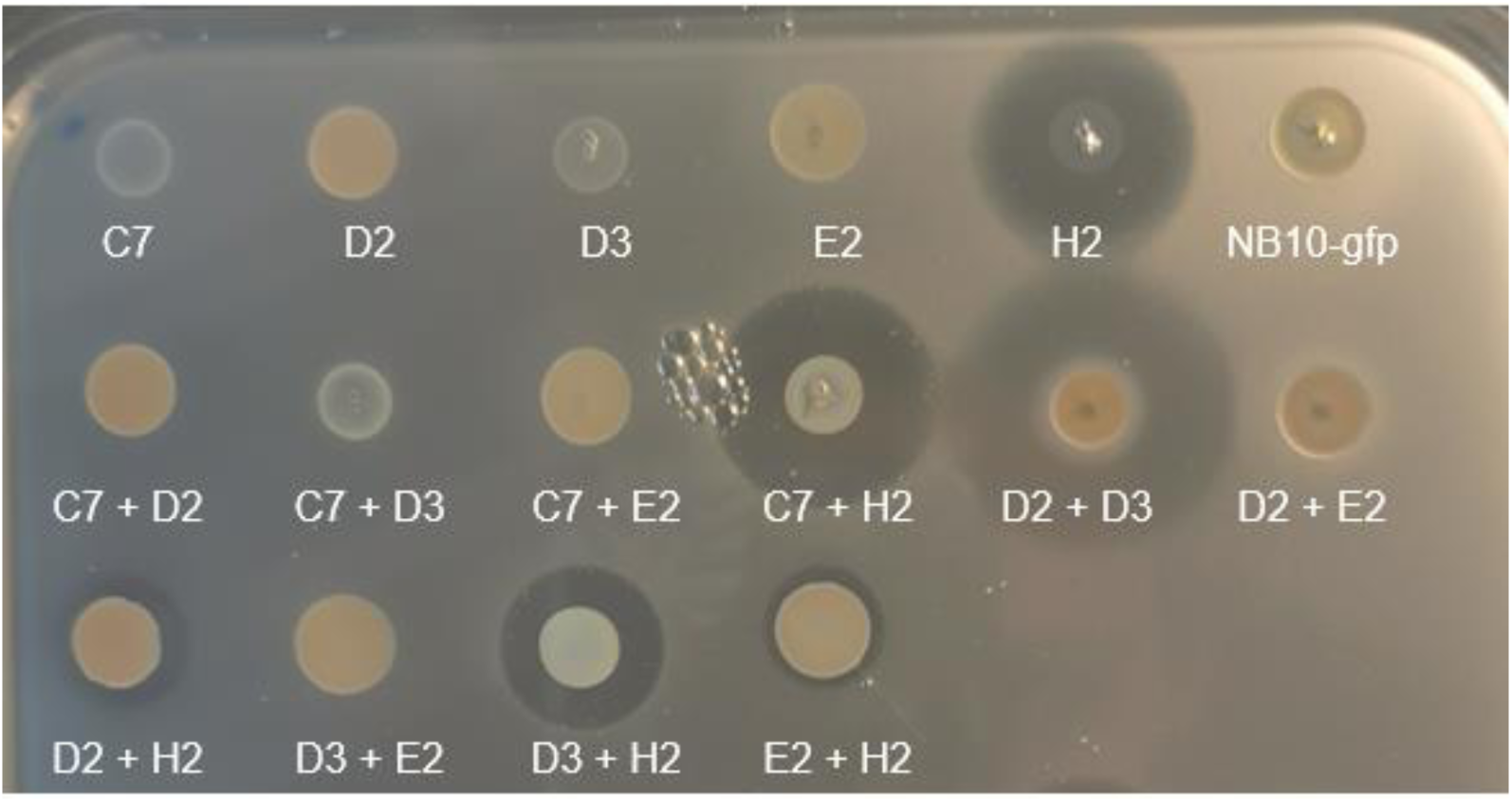
Inhibition of *Vibrio anguillarum* NB10_gfp by co-cultures of bacterial species isolated from the enriched inhibitory *Isochrysis galbana* cultures, in an agar-based pathogen embedding assay. C7: *Roseovarius* sp., D2: *Halomonas campaniensis*, D3: *Sulfitobacter pontiacus*, E2: *Alteromonas macleodii*, H2: *Phaeobacter piscinae*.

**Figure 5.**
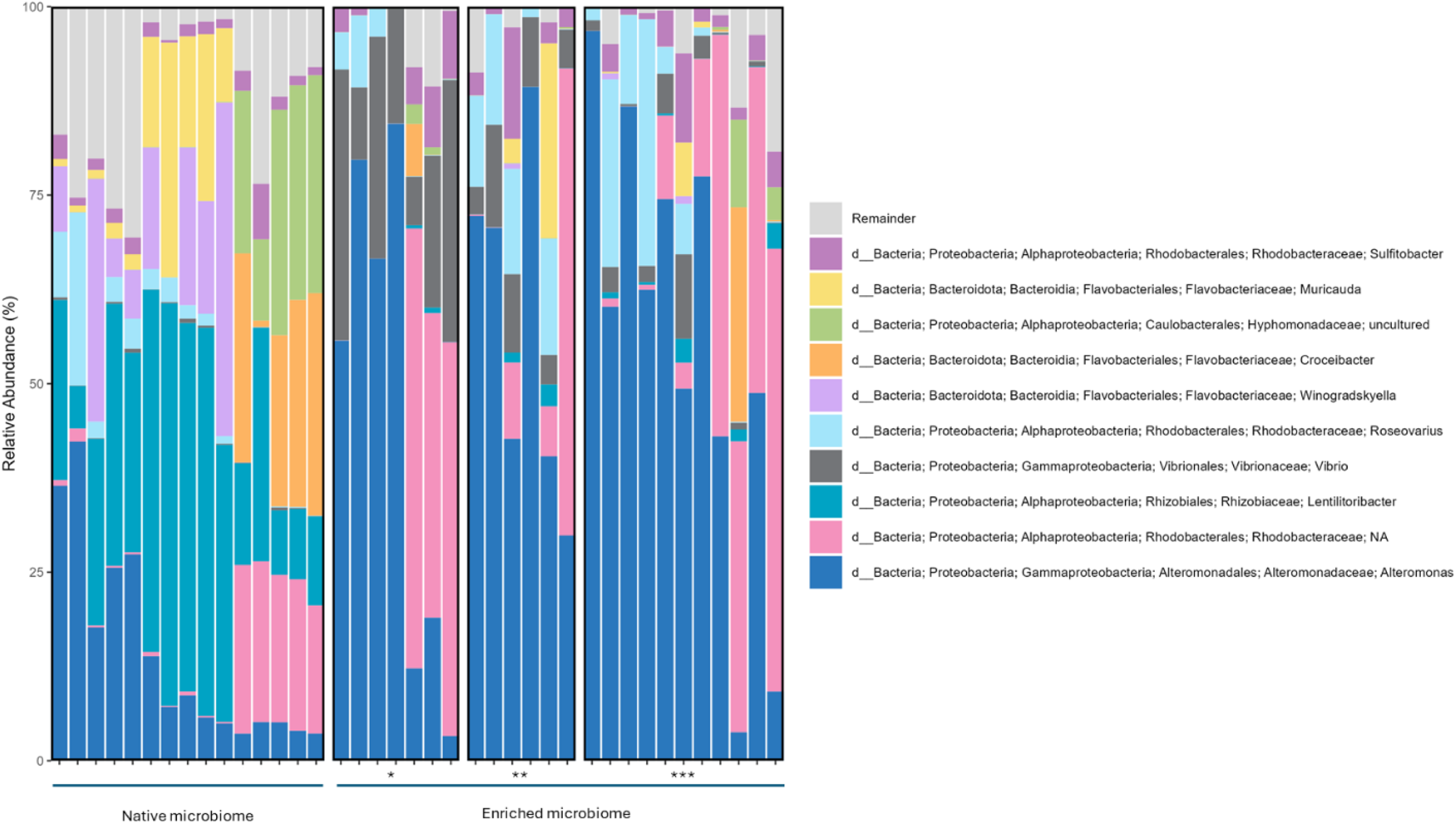
Microbial composition (top 10 most abundant genera) of the native (left) and enriched inhibitory (right) microbiomes based on the 16S rRNA amplicon sequencing results. Samples from the enriched inhibitory microbiome are grouped by their inhibition profile: marginal growth of pathogen (*), medium inhibition of pathogen (**), full inhibition of pathogen (***).

**Table 3.** Bacterial strains isolated from *Isochrysis galbana* cultures with the ability to inhibit *V. anguillarum*. All isolates characterized by a clearing zone of different sizes (+, ++, +++) or a faint clearing zone (*) were identified as *Phaeobacter* sp.

| Isolate | Source | Best BLAST hit | ID% | Inhibition of *Vibrio anguillarum* NB10 |
| --- | --- | --- | --- | --- |
| B7 | Raw culture, NI | *Phaeobacter piscinae* | 99.57% | ++ |
| B9 | Raw culture, NI | *Phaeobacter piscinae* | 99.43% | + |
| C3 | FC, NNI | *Phaeobacter piscinae* | 99.43% | + |
| D4 | FC, NNI | *Phaeobacter piscinae* | 99.14% | + |
| D6 | FC, NI | *Phaeobacter piscinae* | 99.24% | +++ |
| E4 | FC, NI | *Phaeobacter piscinae* | 99.42% | +++ |
| E6 | FC, NI | *Phaeobacter* sp. M8-4.3 | 98.24% | +++ |
| E8 | FC, NI | *Phaeobacter piscinae* | 99.42% | ++ |
| E9 | FC, NI | *Phaeobacter piscinae* | 98.90% | +++ |
| F3 | FC, NI | *Phaeobacter* sp. M8-4.3 | 98.07% | ++ |
| F6 | FM, NNI | *Phaeobacter piscinae* | 98.72% | + |
| H2 | FM, NI | *Phaeobacter piscinae* | 98.96% | +++ |
| H3 | FM, NI | *Phaeobacter piscinae* | 99.33% | * |
| H8 | FM, NI | *Phaeobacter piscinae* | 99.33% | ++ |
| I1 | FM, NI | *Phaeobacter piscinae* | 99.14% | + |
| I2 | FM, NI | *Phaeobacter piscinae* | 99.04% | ++ |
| I3 | FM, NI | *Phaeobacter piscinae* | 99.24% | +++ |

### Whole genome sequencing and genome mining

The species identity of the five chosen isolates was confirmed through Average Nucleotide Identity (ANI%) analysis: *Roseovarius nubinhibens* (isolate C7, ANI 90.0%; referred to as *Roseovarius* sp.), *Halomonas campaniensis* (isolate D2, ANI 98.4%), *Sulfitobacter pontiacus* (isolate D3, ANI 97.3%), *Alteromonas macleodii* (isolate E2, ANI 99.0%), *Phaeobacter piscinae* (isolate H2, ANI 98.2%). Genome assembly of the sequencing data from the five isolates yielded three closed genomes and two draft genomes (Table 4). According to antiSMASH (57) analysis, Halomonas *campaniensis* D2 carries seven BGCs (NI-siderophore, ranthipeptide, redox-cofactor, RIPP-like, T1PKS, betalactone, ectoine) and *Sulfitobacter pontiacus* D3 four BGCs (RiPP-like, betalactone, hserlactone, redox-cofactor), potentially encoding antibacterial compounds (Table S6). The structure of compounds cannot be accurately predicted due to the low similarity compared to the known BGCs in MIBiG (Table S6).

**Table 4.**
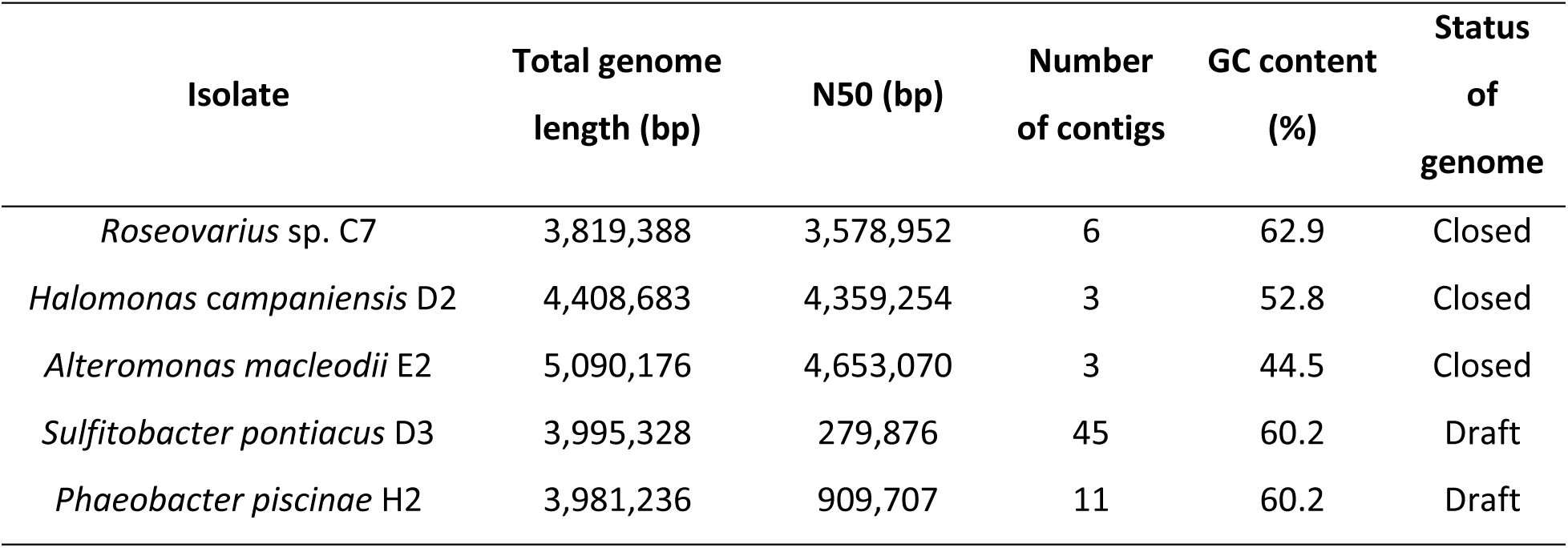
Whole genome sequencing results of five bacterial isolates from the *Isochrysis galbana* microbiome. Whole genome sequencing of *Roseovarius* sp. C7, *Halomonas campaniensis* D2, *Sulfitobacter pontiacus* D3, *Alteromonas macleodii* E2, and *Phaeobacter piscinae* H2 yielded three closed genomes and two draft genomes.

| Isolate | Total genome length (bp) | N50 (bp) | Number of contigs | GC content (%) | Status of genome |
| --- | --- | --- | --- | --- | --- |
| *Roseovarius* sp. C7 | 3,819,388 | 3,578,952 | 6 | 62.9 | Closed |
| *Halomonas campaniensis* D2 | 4,408,683 | 4,359,254 | 3 | 52.8 | Closed |
| *Alteromonas macleodii* E2 | 5,090,176 | 4,653,070 | 3 | 44.5 | Closed |
| *Sulfitobacter pontiacus* D3 | 3,995,328 | 279,876 | 45 | 60.2 | Draft |
| *Phaeobacter piscinae* H2 | 3,981,236 | 909,707 | 11 | 60.2 | Draft |

## DISCUSSION

The rapid spread of pathogenic bacteria within intensive aquaculture systems remains a significant challenge, especially during the vulnerable larval development stage, and disease outbreaks often result in substantial economic losses. In this study, we investigated the probiotic potential of the microbiomes from two algae typically employed as live feed in marine larviculture, *Tetraselmis suecica* and *Isochrysis galbana*. By identifying microbiomes with disease suppression, we aim to provide a novel approach to disease control in aquaculture, reducing reliance on antibiotics and the risk of spread of AMR.

Determining the inhibitory effect of a mixed microbiome against a particular pathogenic bacterium requires specific detection and quantification of the pathogen. Previous works on fish pathogens have used several approaches for determining inhibition in a mixed microbial population. For instance tagging the pathogen with an antibiotic resistance marker enabling specific quantification by culturing on antibiotic containing media (11, 20, 31), using semi-selective media and distinct colony morphology in defined co-cultures (8, 59, 60), using flow cytometry to determine concentrations of a GFP-tagged pathogen (20), or developing specific primers and designing a qPCR approach has been a strategy for specific detection (8). Both the culturing approaches as well as the qPCR method are labour-intensive and time-consuming. To enable a faster screen for pathogen-inhibitory effect of a mixed microbial community, we used a GFP-tagged pathogen and demonstrated that the fluorescent signal is a robust indication of growth of the pathogen, and that it can be used to detect its inhibition by other microorganisms. Thus, we developed a simple methodology for screening mixed microbiomes for their anti-pathogenic potential, thereby offering a valuable tool for developing sustainable disease management strategies also outside of the aquaculture sector. The method facilitates identification and selection of microbiomes with the greatest potential for disease control, contributing to the sustainable growth of aquaculture and the global effort to combat AMR.

We used the method to demonstrate that both the native, complete microbiomes and diluted microbiomes of two microalgae used in aquacultures were able to inhibit the growth of fish pathogen *Vibrio anguillarum*. In contrast to findings from previous studies where cultures of *Isochrysis* sp. and *Tetraselmis suecica* have themselves been inhibitory to fish pathogenic *Vibrio* species (59–61), we did not find that *V. anguillarum* was inhibited by axenic *I. galbana* or *T. suecica,* nor the cell-free supernatants of either culture. Thus, in our study, it was clearly the microorganisms associated with the algae that inhibited growth of the pathogen, emphasising the crucial role of microbial communities associated with live feed in larviculture for disease control. These findings highlight the potential of harnessing natural microbial consortia to enhance disease resistance in aquaculture settings. Furthermore, in our experiments the *Isochrysis* microbiome inhibited growth of the pathogen (RFU < 10% of control) at 1:10 (*Vibrio*: *Isochrysis* microbiome) starting inoculation ratio, showing a more pronounced inhibitory potential as compared to that of a monoculture of *P. piscinae* S26, where it took an inoculation ratio of 1:100 (*Vibrio*: *P. piscinae* S26) to reach similar reduction of the RFU, supporting our hypothesis that a mixed microbiome may be more inhibitory to a pathogen than individual pure cultures.

Sequence-based analysis of *I. galbana* microbiomes found that *Rhizobiaceae*, *Alteromonadaceae*, and *Flavobacteriaceae* family members dominated the native *I. galbana* microbiome. This is in line with a number of previous studies reporting these families as members often found in the core microbiome of microalgae (36, 62, 63). The native microbiome also harboured members of the *Rhodobacteraceae*, which is a family commonly associated with microalgae and algal blooms (36, 62–65). The enriched inhibitory microbiomes were dominated by *Alteromonadaceae*, *Rhodobacteraceae*, and *Vibrionaceae* members, with decreased relative abundances of *Rhizobiaceae* and *Flavobacteriaceae* compared to the native microbiome. These shifts are likely heavily influenced by transferring the algal microbiome to a nutrient rich assay medium (MB) favouring fast-growing, heterotrophic bacteria. Interestingly, the degree of pathogen inhibition by the enriched inhibitory microbiomes is not explained by differences in the community richness, diversity, and evenness. However, it does correlate with distinct patterns in the overall community composition, suggesting that the inhibitory potential is influenced by the structure of the microbiome and synergistic interactions within a diverse community. Our findings also imply that complete pathogen inhibition can be achieved by a range of different enrichment cultures.

Pure cultures of *Phaeobacter piscinae* H2 isolated from the *I. galbana* microbiome inhibited *V. anguillarum* when tested in an agar-based assay, and this is consistent with numerous previous studies showing that several *Phaeobacter* species can inhibit *Vibrio* species (20, 21, 66–68). Notably, all isolates that produced an inhibition zone of the embedded *V. anguillarum* were identified as *Phaeobacter* species, however, this genus was not among the ten most abundant genera of the enriched inhibitory microbiomes. This could indicate that instead of the potent antagonistic effect of particular genera, it may be a synergistic effect of the microbiome members that inhibits the pathogen.

The co-culture of *Sulfitobacter pontiacus* D3 and *Halomonas campaniensis* D2, isolated from the *I. galbana* microbiome, was inhibitory to *V. anguillarum* NB10, despite neither isolate showing pathogen inhibition as pure cultures. The co-culture also showed varying degrees of inhibition against three additional *V. anguillarum* strains, belonging to different phylogenetic clades based on MLSA, including a faint inhibitory effect of the highly virulent *V. anguillarum* PF7. This indicates that the observed inhibition is not limited to low-virulence strains but also that the effect may not be universal and that some pathogens are not affected. While it is beyond the scope of the current study to unravel the molecular and chemical mechanism underlying this co-culture-induced inhibition, previous research has demonstrated that biosynthetic gene clusters (BGCs) encoding antibiotic compounds can be elicited by co-cultivation of bacteria (28–30, 69). According to antiSMASH analyses, the genomes of both bacteria harbour several BGCs. Therefore, future metabolomic analyses of the mono- and co-cultures could provide insight into the co-culture-induced antimicrobial mechanism. Moreover, several *Halomonas* species have been reported to produce an array of hydrolytic enzymes, extracellular polysaccharides (EPS) (70), antibiotics (71), antifungals (72), and pigments (73). *Halomonas campaniensis* D2, in particular, carries seven biosynthetic gene clusters potentially encoding antibacterial compounds. This combined with the lack of inhibition by the monoculture could suggest that the presence of *Sulfitobacter pontiacus* D3 may trigger the inhibitory effect. *Sulfitobacter* species, often associated with marine algae and corals (74–76), possess the ability to metabolise sulphite (77–79) and dimethylsulfoniopropionate (DMSP) (74). The breakdown of DMSP produces potentially antimicrobial compounds such as acrylate (80, 81) and dimethyl sulphide (DMS), which could indicate that the sulphur metabolism capabilities of *Sulfitobacter* may contribute to pathogen inhibition. Several *Sulfitobacter* species have also been reported to inhibit fungal pathogens (82) and produce EPS (83) and extracellular cyclodipeptides (84).

Beiralas et. al reported that *Sulfitobacter pontiacus* is a key species of the *Emiliania huxleyi* microbiome, protecting its algal host from bacterial pathogens even at low initial abundance (76). While the mechanism of the algal protector phenotype was not determined, the authors suggest that harbouring *Sulfitobacter* species in the algal microbiome presents the host with an advantage and that *Sulfitobacter* species may play a key role in influencing population dynamics. Building on these findings, we speculate that even a low abundance of *Sulfitobacter pontiacus* in a community or co-culture may induce synergistic effects that enhance pathogen inhibition, potentially mediated through the production of bioactive compounds. These insights highlight the importance of microbial interactions in community functionality and stability.

In conclusion, by introducing an efficient methodology for screening mixed microbiomes, our study offers a promising approach for identifying candidate microbiomes for sustainable disease management. Our findings highlight the potential of harnessing natural microbial consortia to enhance disease resistance in aquaculture and support the importance of microbial co-cultures as potential disease control strategies.

## ACKNOWLEDGEMENTS

We thank our aquaculture partners for providing algal cultures. This project has received funding from the European Union’s Horizon 2020 research and innovation programme under Grant Agreement no. 101000392 (MARBLES), the Novo Nordisk Foundation (NNF20OC0064249) and the Danish National Research Foundation (DNRF137) for the Center for Microbial Secondary Metabolites. This output reflects only the author’s view, and the Research Executive Agency (REA) cannot be held responsible for any use that may be made of the information contained therein. Figure 1 was created with BioRender.com.

## Supplementary Tables and Figures

**Table S1.**
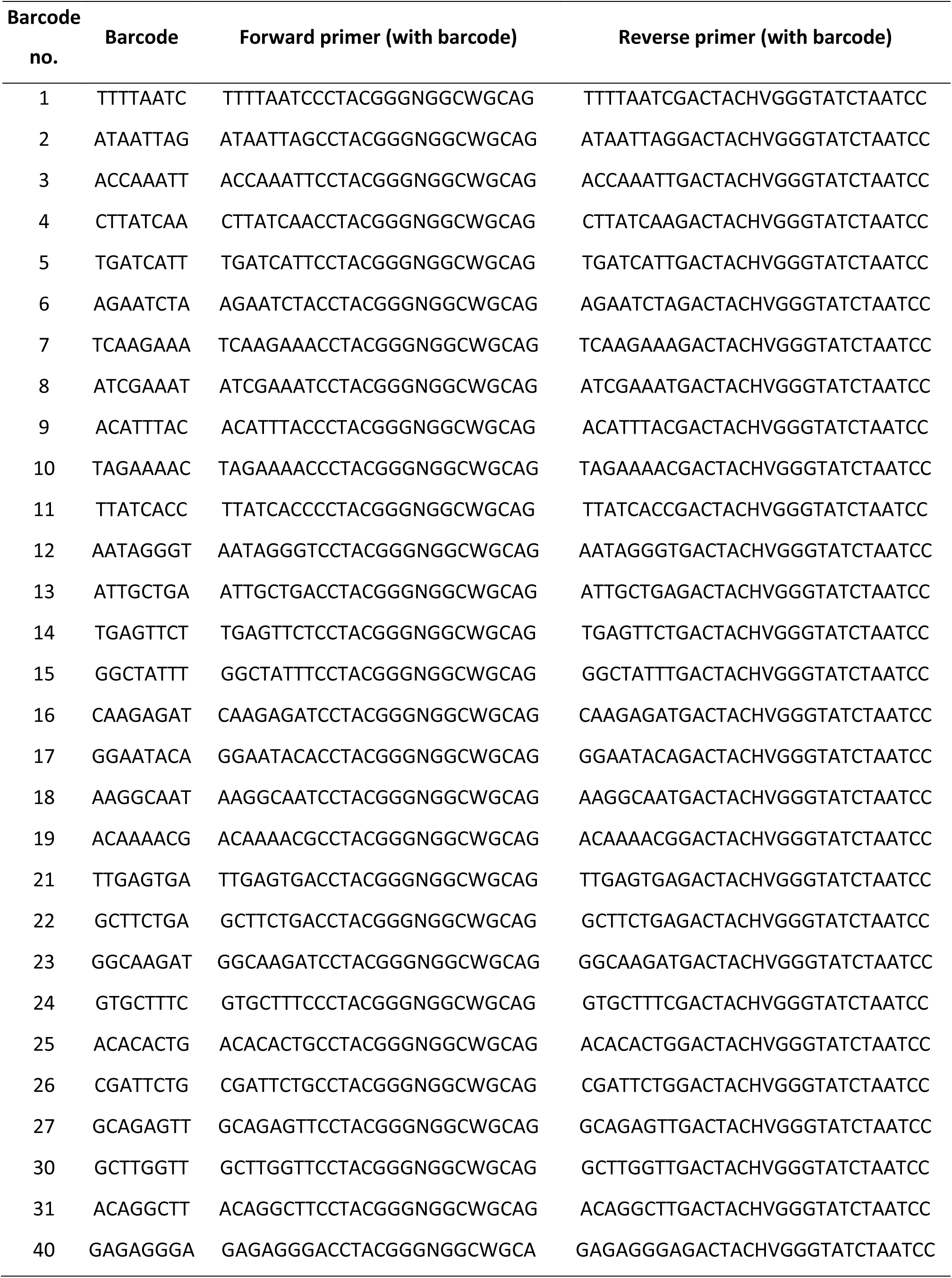
List of thirty unique barcodes and primers used to amplify the V3-V4 region of the 16S rRNA gene in DNA from the microbiomes of *Isochrysis galbana* cultures. Primers and barcodes from(85).

**Table S2.**
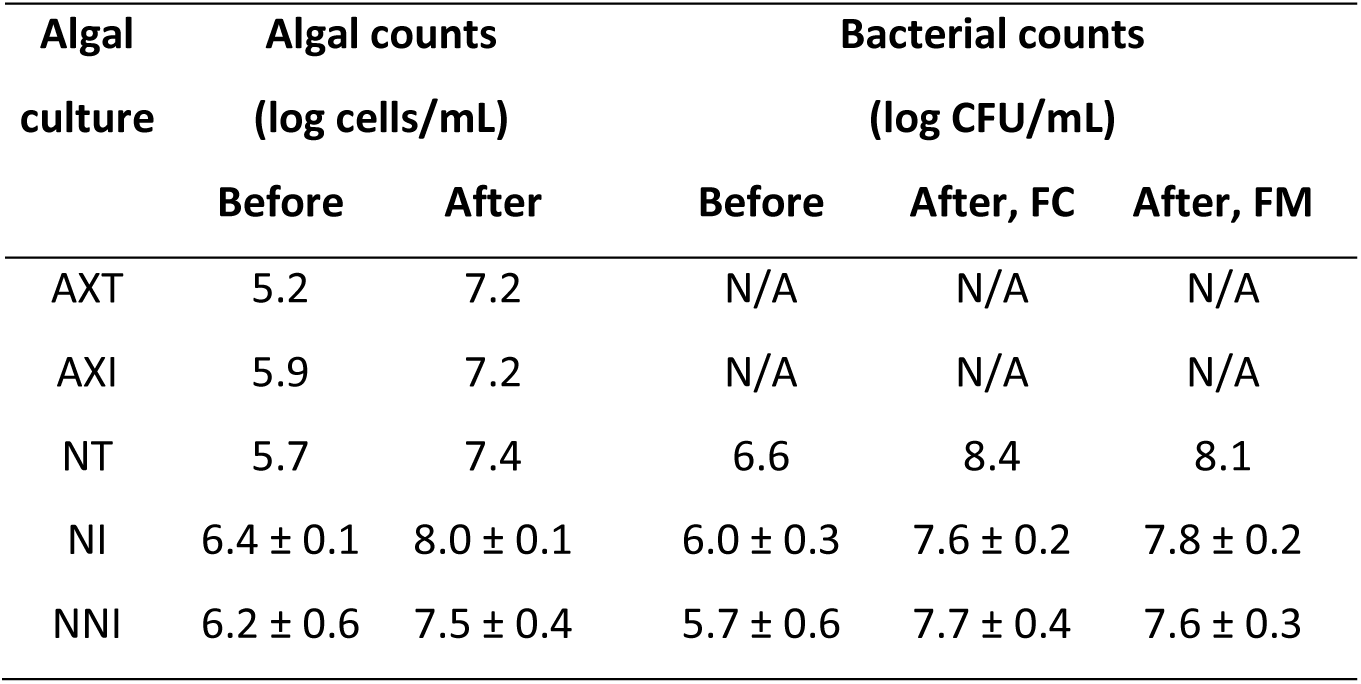
Algal and bacterial counts of different algal cultures (AXT, AXI, NT, NI and NNI) used in the *Vibrio* inhibition assay, before and after up-concentration (“Before” and “After”), and from different algal sample fractions (Full culture, FC and Filtered microbiome, FM). Counts are reported as mean ± standard deviation. In cases where standard deviation is not reported, data originated from unreplicated experimental conditions.

| Algal culture | Algal counts<br>(log cells/mL) |  | Bacterial counts<br>(log CFU/mL) |  |  |
| --- | --- | --- | --- | --- | --- |
|  | Before | After | Before | After, FC | After, FM |
| AXT | 5.2 | 7.2 | N/A | N/A | N/A |
| AXI | 5.9 | 7.2 | N/A | N/A | N/A |
| NT | 5.7 | 7.4 | 6.6 | 8.4 | 8.1 |
| NI | 6.4 ± 0.1 | 8.0 ± 0.1 | 6.0 ± 0.3 | 7.6 ± 0.2 | 7.8 ± 0.2 |
| NNI | 6.2 ± 0.6 | 7.5 ± 0.4 | 5.7 ± 0.6 | 7.7 ± 0.4 | 7.6 ± 0.3 |

**Table S3.**
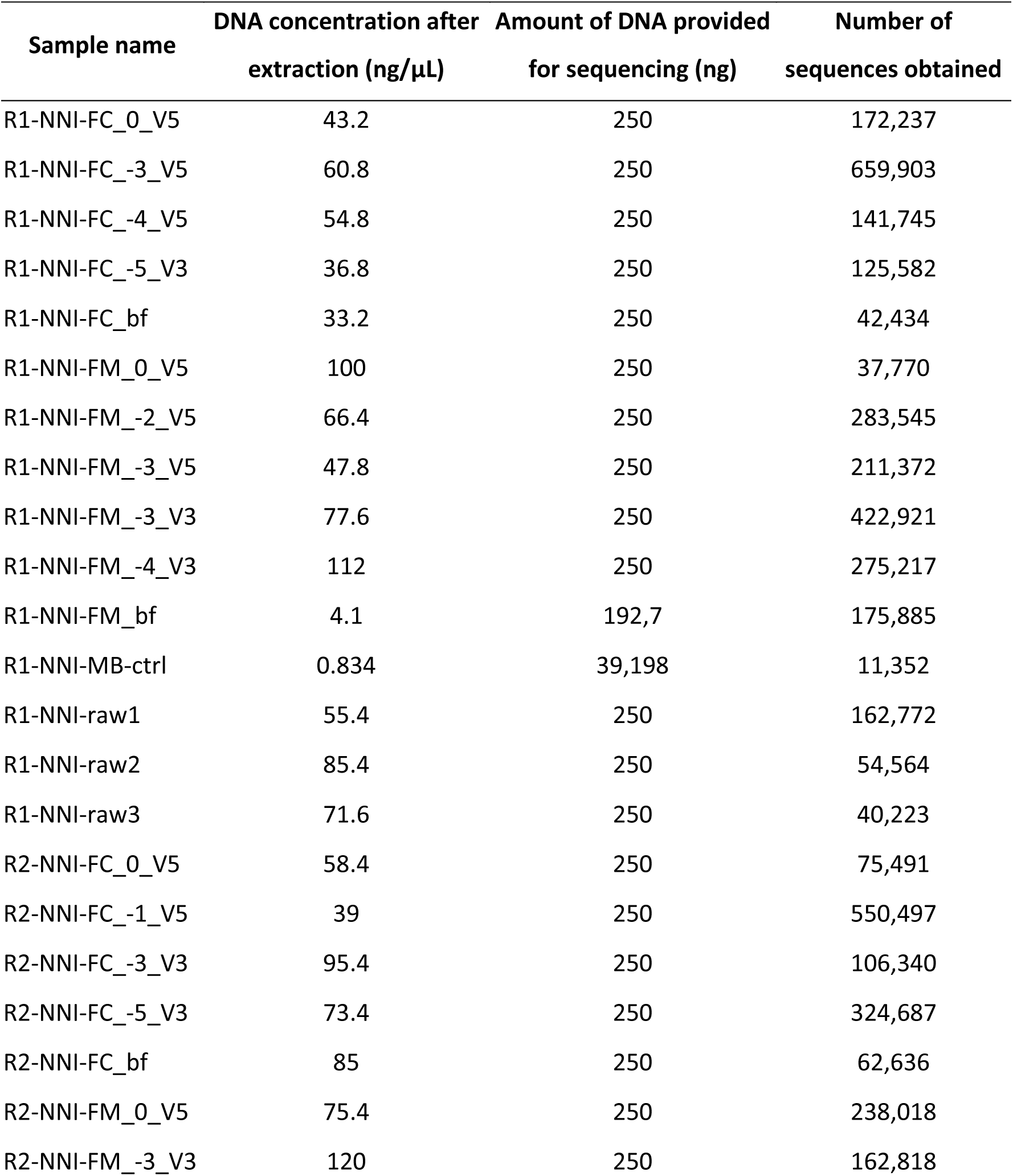
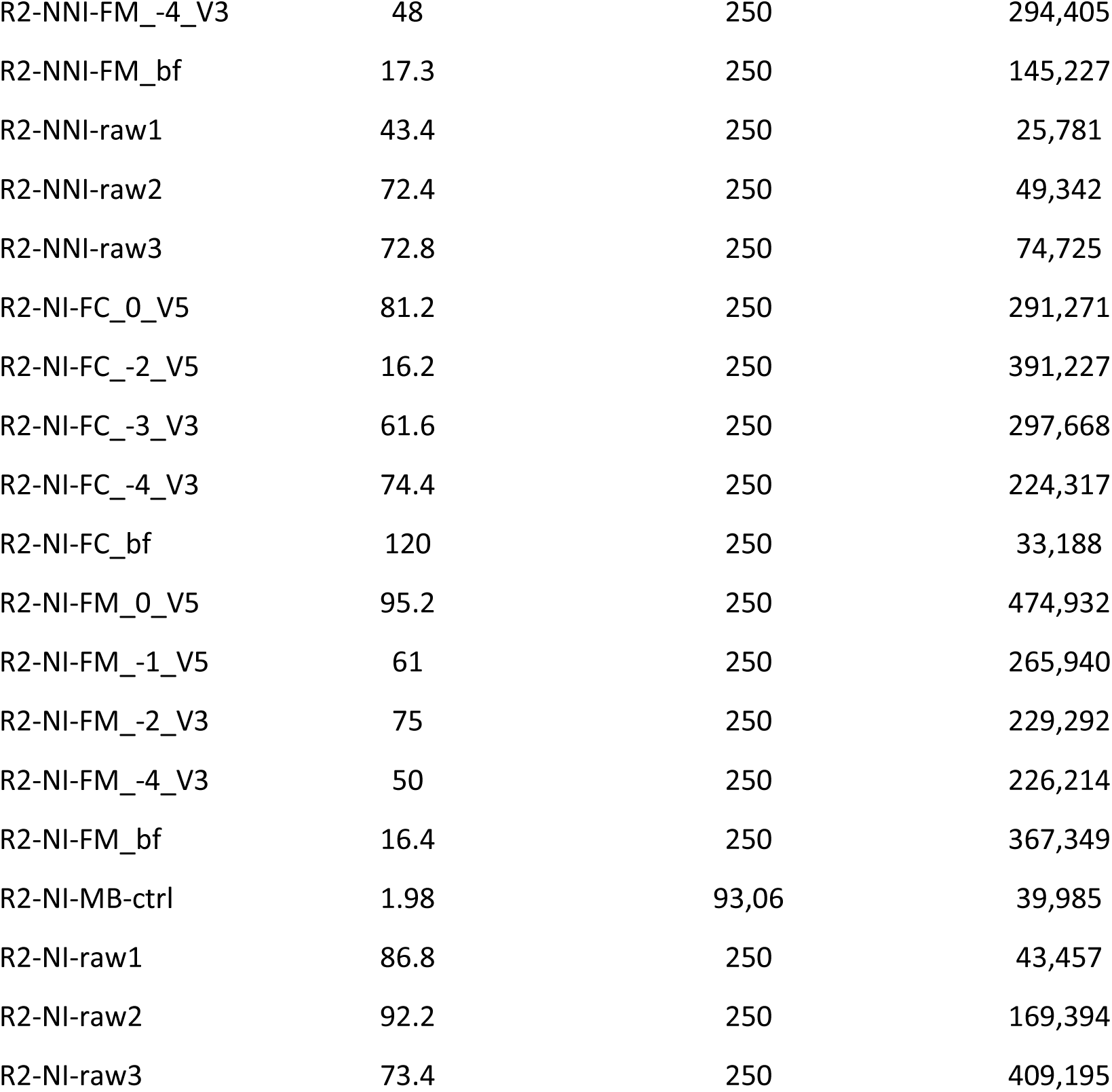
DNA concentration after extraction, DNA amount provided for sequencing and number of sequences obtained from each sample (after denoising and filtering). Sample names indicate replicate (“R1”, “R2”) source culture (“NI”, “NNI”), sample fraction (Full culture: “FC”, Filtered microbiome: “FM”), dilution factor of the algal microbiome (0 to −5), and starting inoculum of *V. anguillarum* NB10_gfp (“V3”, “V5” for 3 or 5 log CFU mL^-1^) for samples obtained after the inhibition assay. The algal cultures were sampled before up-concentrating and filtering (“raw”, triplicates), and each fraction (FC and FM) was sampled before the enrichment assay (“bf”). Sterile media controls were included (“MB-ctrl”).

**Table S4.**
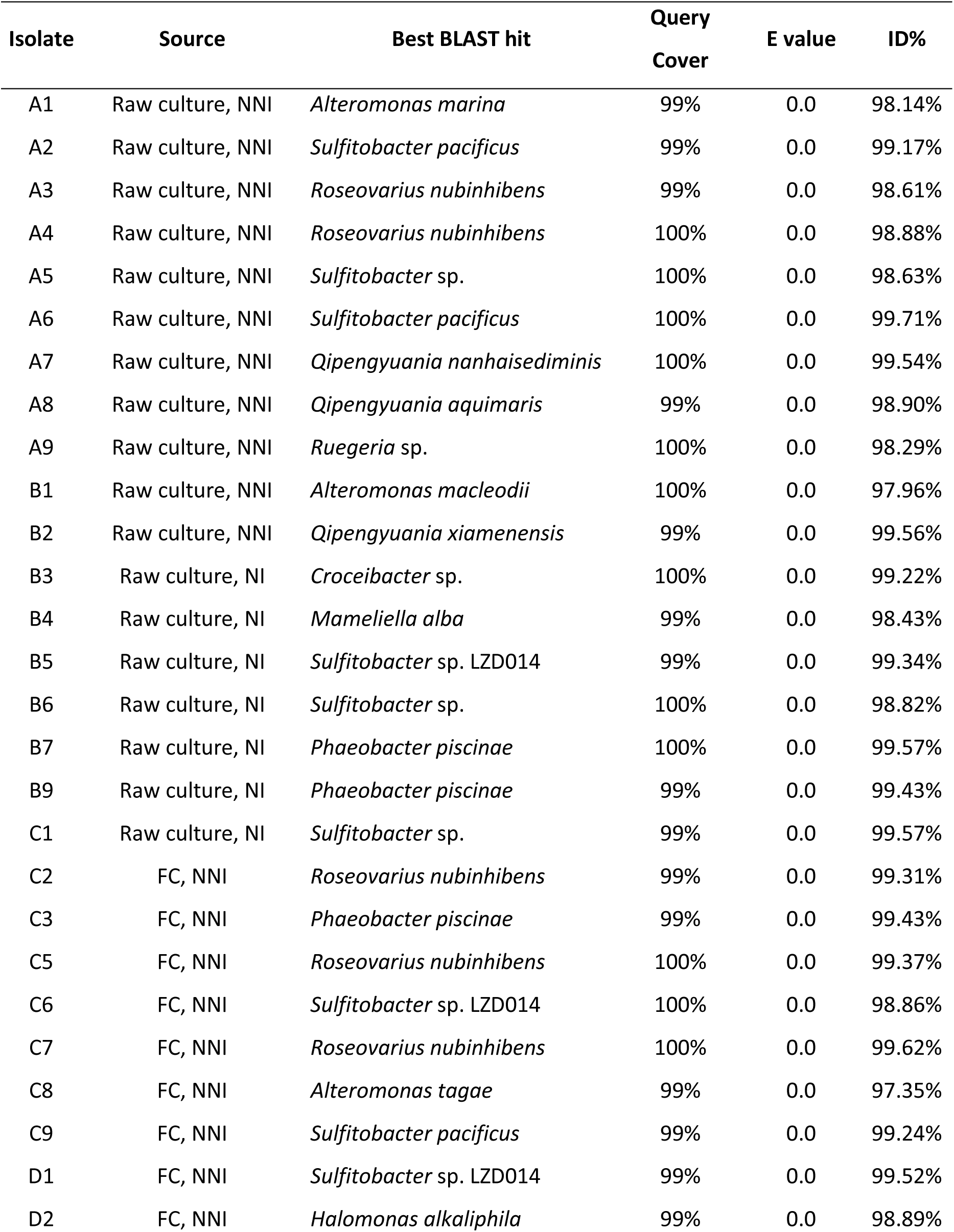
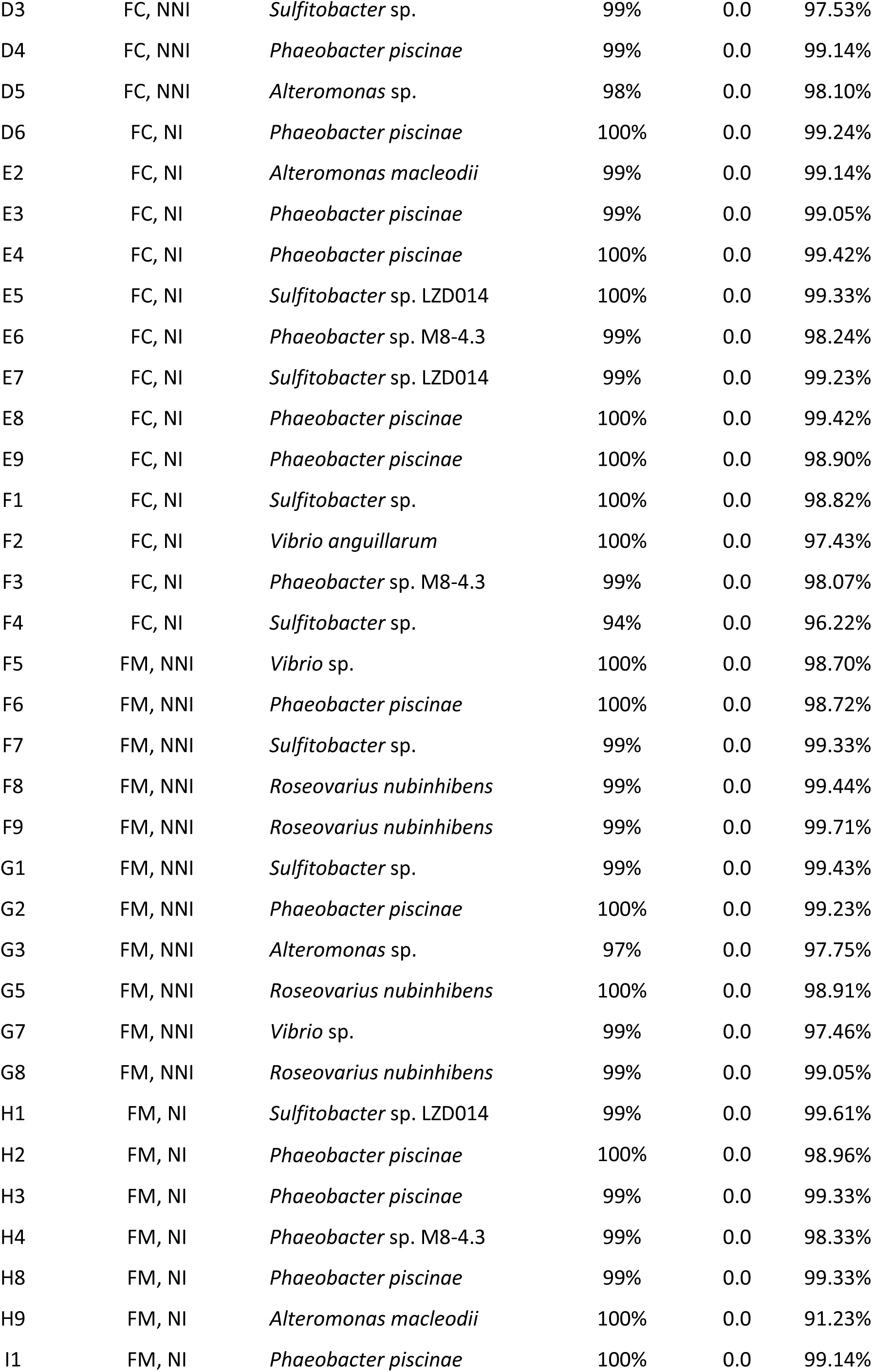
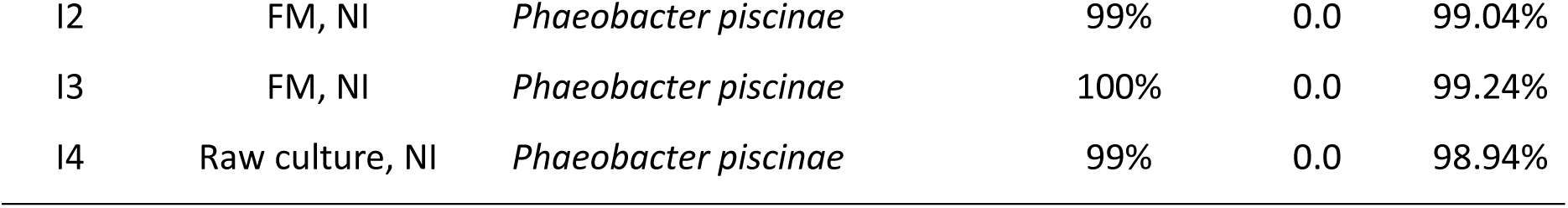
64 bacterial strains isolated from the native („Raw culture”, 19 strains) or enriched inhibitory („Full culture”: FC, or „Filtered microbiome”: FM, 45 strains) microbiome of *Isochrysis galbana*, and tentatively identified using full-length 16S rRNA gene sequencing. The isolates originate from two different *I. galbana* cultures (NI and NNI), acquired at different times from the same aquaculture unit.

**Table S5.**
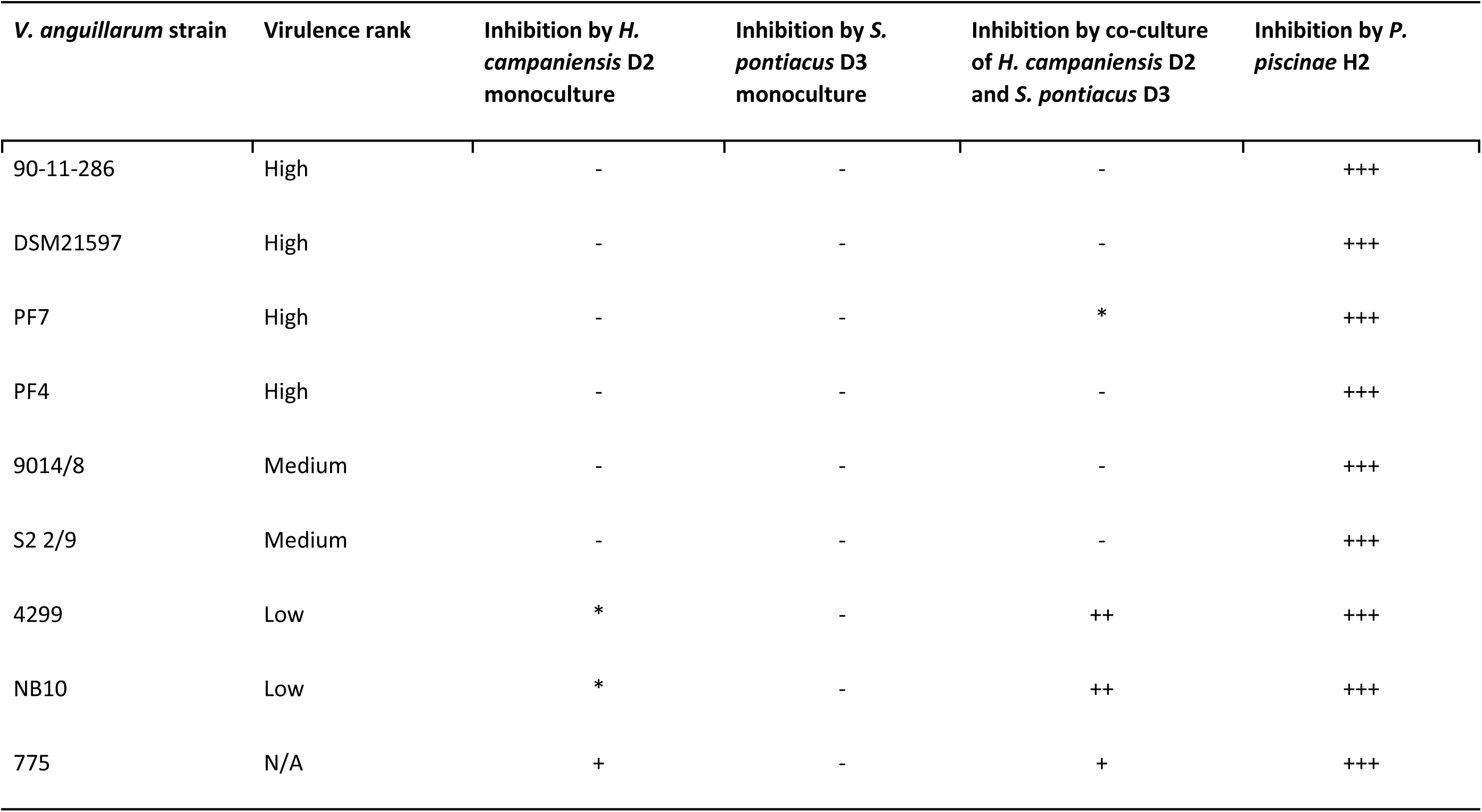
Inhibition of nine *V. anguillarum* strains by the co-culture of *Halomonas campaniensis* (isolate D2) and *Sulfitobacter pontiacus* (isolate D3). *Phaeobacter piscinae* isolate H2 was used as positive control. Clearing zone of different degrees (+, ++, +++), a faint clearing zone (*), or no clearing zone (-) after 24 hours of incubation. Virulence ranks as defined by Rønneseth et al (54).

**Table S6.**
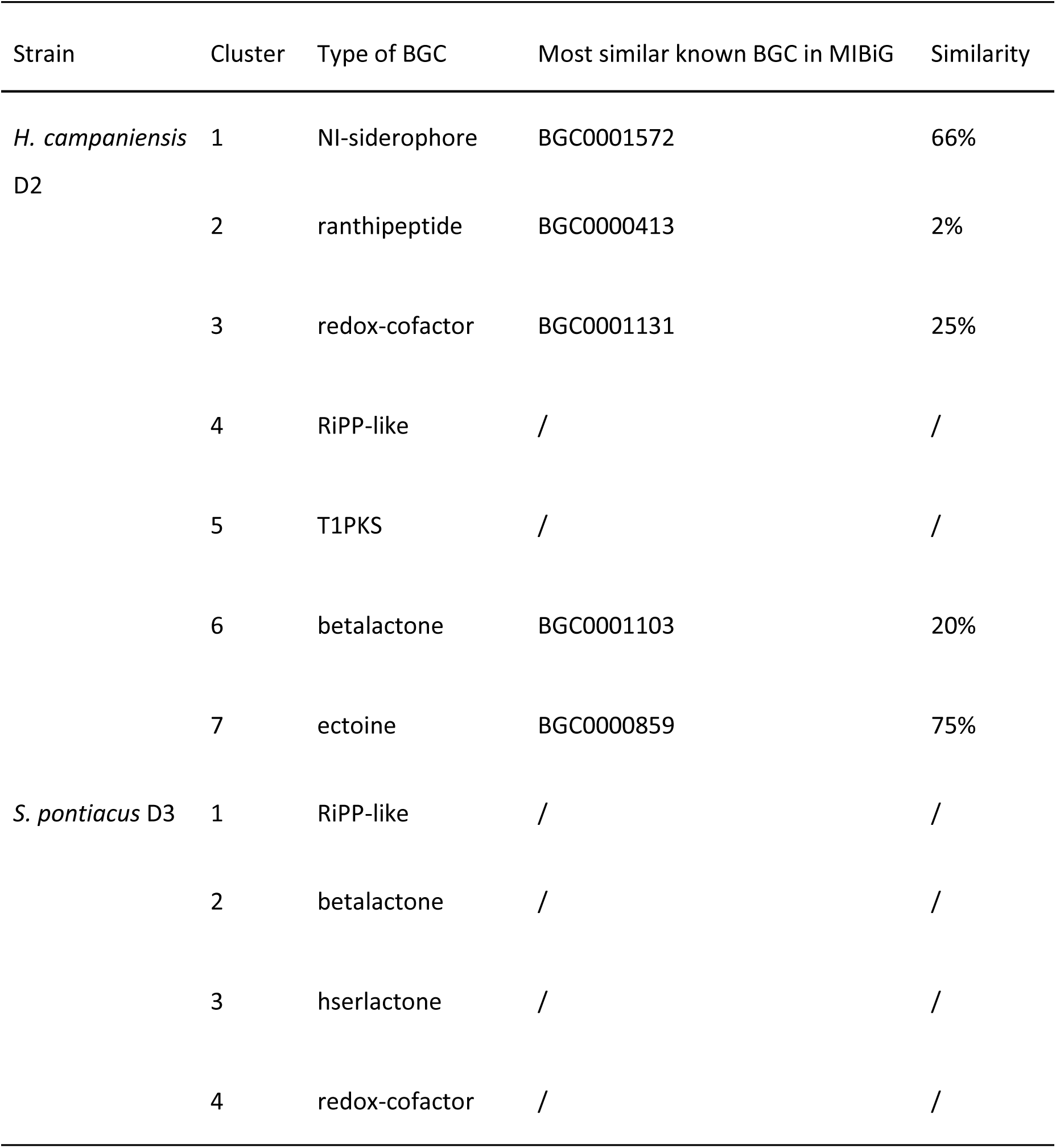
Biosynthetic gene clusters (BGCs) predicted by antiSMASH in the genomes of *Halomonas campaniensis* D2 and *Sulfitobacter pontiacus* D3.

**Figure S1.**
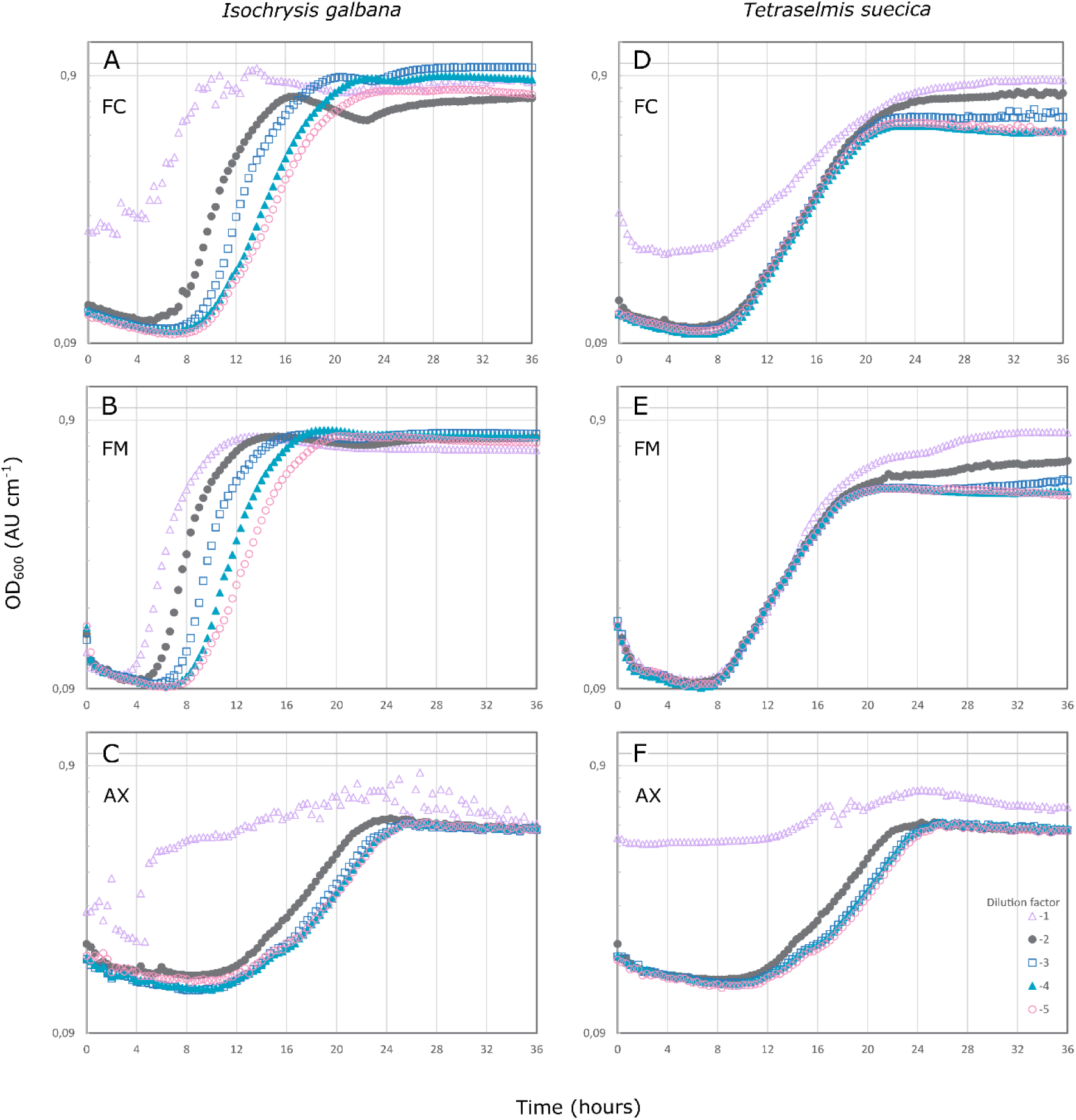
Inhibition assay by *Isochrysis galbana* (left) and *Tetraselmis suecica* (right) microbiomes against *Vibrio anguillarum* NB10_gfp, at a starting concentration of 3.1 ± 0.3 log CFU mL^-1^, as measured by absorbance at 600 nm. The inhibitory effect of serial dilutions (10^-1^ 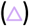, 10^-2^ 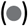, 10^-3^ 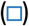, 10^-4^ 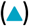, 10^-5^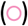) of different fractions of the algal cultures have been tested: full culture (FC; S1A and S1D), with algal and bacterial cells; filtered microbiome (FM; S1B and S1E), where algal cells have been removed; and axenic (AX; S1C and S1F) cultures, where the algae cells are free of bacteria.

**Figure S2.**
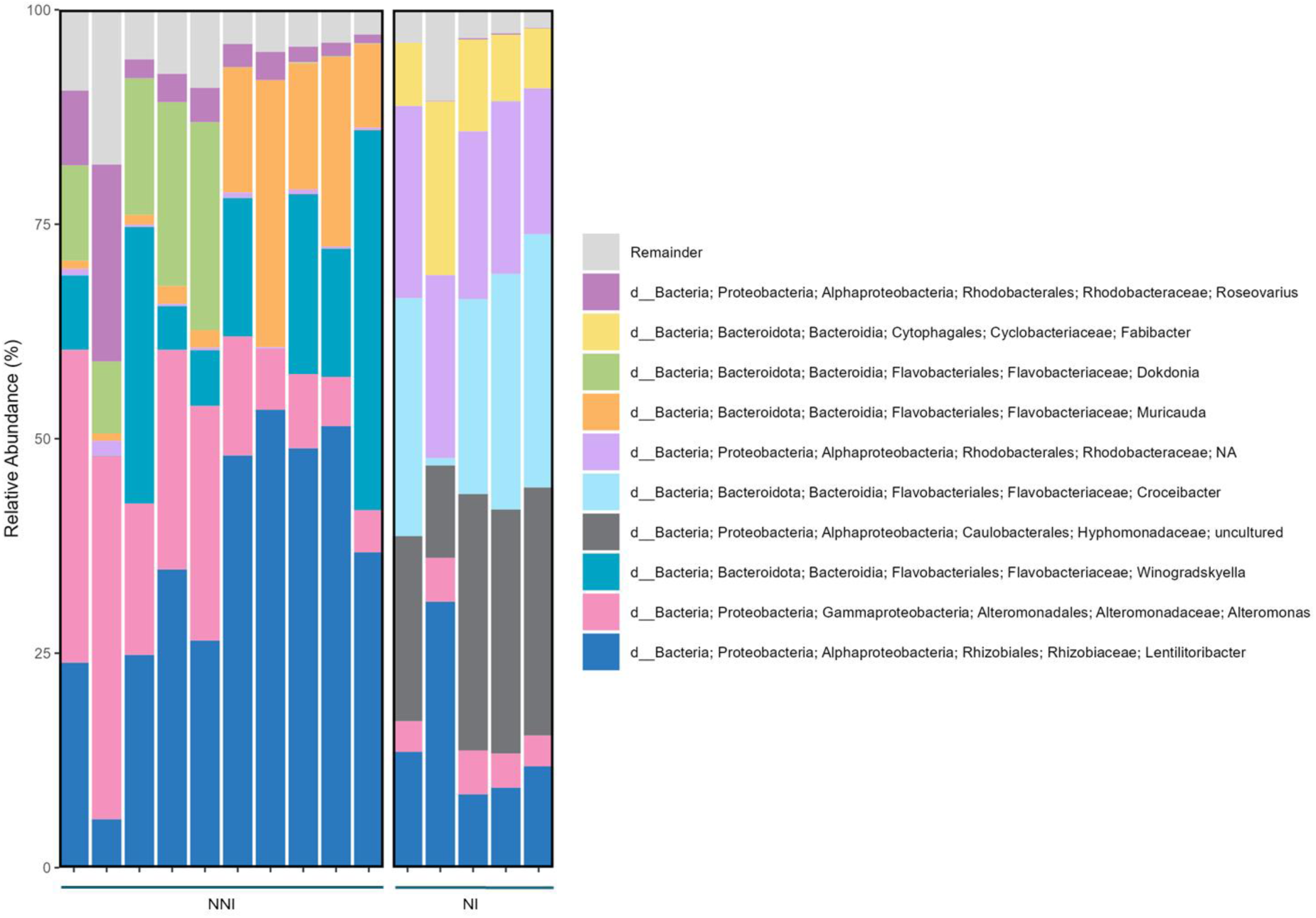
Native microbiome composition (top 10 most abundant genera) of two *Isochrysis galbana* cultures of different age based on the 16S rRNA amplicon sequencing results. One culture, NNI has been newly provided by an aquaculture facility and has a relatively freshly recruited microbiome (left). The other culture, NI has been regularly subcultured under laboratory conditions for almost four years (right).

**Figure S3.**
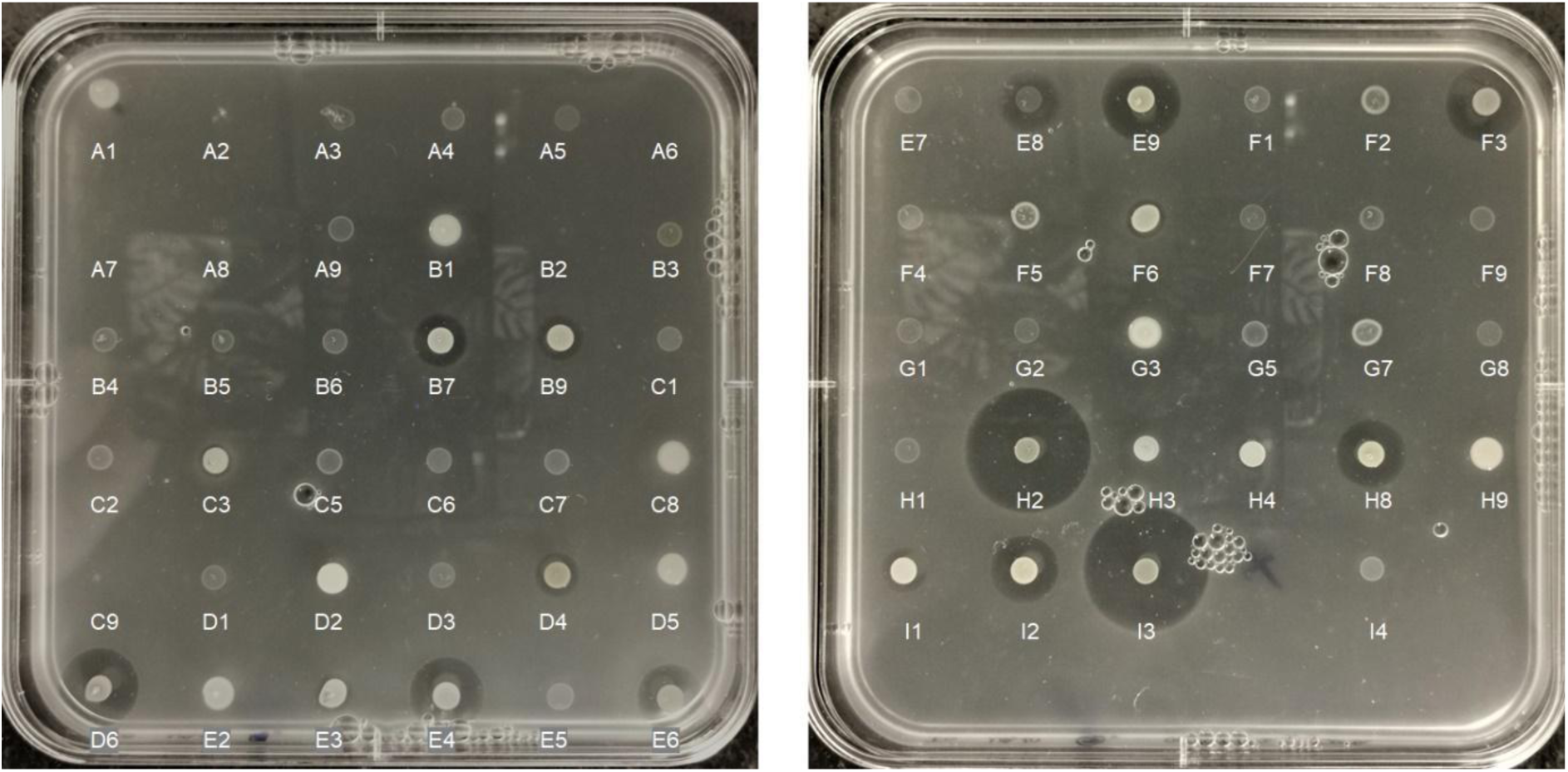
Inhibition of *Vibrio anguillarum* NB10_gfp by 64 isolates from the *Isochrysis galbana* microbiome. 16 isolates showed a clearing zone of varying size (all identified as *Phaeobacter* sp.), and 13 isolates showed a faint halo, or a rough agar surface around the colony (identified as *Alteromonas, Sulfitobacter, Qipengyuania* and *Croceibacter* sp.).

## Notes

### Competing Interest Statement

The authors have declared no competing interest.

### Summary of Updates

New experiments added, testing against several V. anguillarum strains. Also, determining the # of vibrio in the gfp assay. Introduction rewritten

## REFERENCES

1. FAO. 2022. The State of World Fisheries and Aquaculture 2022. Towards Blue Transformation. FAO; https://openknowledge.fao.org/handle/20.500.14283/cc0461en.

2. Ringø E, Olsen RE, Jensen I, Romero J, Lauzon HL. 2014. Application of vaccines and dietary supplements in aquaculture: possibilities and challenges. Rev Fish Biol Fisheries 24:1005–1032.

3. Westerdahl A, Olsson JC, Kjelleberg S, Conway PL. 1991. Isolation and characterization of turbot (*Scophtalmus maximus*)-associated bacteria with inhibitory effects against *Vibrio anguillarum*. Applied and Environmental Microbiology 57:2223–2228.

4. Ina-Salwany MY, Al-Saari N, Mohamad A, Mursidi F-A, Mohd-Aris A, Amal MNA, Kasai H, Mino S, Sawabe T, Zamri-Saad M. 2019. Vibriosis in Fish: A Review on Disease Development and Prevention. J Aquat Anim Health 31:3–22.

5. Frans I, Michiels CW, Bossier P, Willems KA, Lievens B, Rediers H. 2011. *Vibrio anguillarum* as a fish pathogen: virulence factors, diagnosis and prevention: Pathogen profile of *Vibrio anguillarum*. Journal of Fish Diseases 34:643–661.

6. Mohamad N, Amal MNA, Yasin ISM, Zamri Saad M, Nasruddin NS, Al-saari N, Mino S, Sawabe T. 2019. Vibriosis in cultured marine fishes: a review. Aquaculture 512:734289.

7. Hamre K, Yufera M, Rønnestad I, Boglione C, Conceição L, Izquierdo M. 2013. Fish larval nutrition and feed formulation: Knowledge gaps and bottlenecks for advances in larval rearing. Reviews in Aquaculture 5:S26–S58.

8. Rasmussen BB, Erner KE, Bentzon-Tilia M, Gram L. 2018. Effect of TDA-producing *Phaeobacter inhibens* on the fish pathogen *Vibrio anguillarum* in non-axenic algae and copepod systems. Microbial Biotechnology 11:1070–1079.

9. Reid HI, Treasurer JW, Adam B, Birkbeck TH. 2009. Analysis of bacterial populations in the gut of developing cod larvae and identification of *Vibrio logei*, *Vibrio anguillarum* and *Vibrio splendidus* as pathogens of cod larvae. Aquaculture 288:36–43.

10. Dittmann KK, Rasmussen BB, Castex M, Gram L, Bentzon-Tilia M. 2017. The aquaculture microbiome at the centre of business creation. Microbial Biotechnology 10:1279–1282.

11. D’Alvise PW, Lillebø S, Prol-Garcia MJ, Wergeland HI, Nielsen KF, Bergh Ø, Gram L. 2012. *Phaeobacter gallaeciensis* reduces *Vibrio anguillarum* in cultures of microalgae and rotifers, and prevents vibriosis in cod larvae. PLoS ONE 7:e43996.

12. Gatesoupe FJ. 1999. The use of probiotics in aquaculture. Aquaculture 180:147–165.

13. Kesarcodi-Watson A, Kaspar H, Lategan MJ, Gibson L. 2008. Probiotics in aquaculture: The need, principles and mechanisms of action and screening processes. Aquaculture 274:1–14.

14. Hotel A. 2001. Health and Nutritional Properties of Probiotics in Food Including Powder Milk with Live Lactic Acid Bacteria – Joint FAO/WHO Expert Consultation 2014.

15. Sonnenschein EC, Jimenez G, Castex M, Gram L. 2021. The *Roseobacter* -Group Bacterium *Phaeobacter* as a Safe Probiotic Solution for Aquaculture. Appl Environ Microbiol 87:e02581–20.

16. Lieke T, Meinelt T, Hoseinifar SH, Pan B, Straus D, Steinberg C. 2019. Sustainable aquaculture requires environmental-friendly treatment strategies for fish diseases. Reviews in Aquaculture 12:943–965.

17. Hoseinifar SH, Sun Y-Z, Wang A, Zhou Z. 2018. Probiotics as Means of Diseases Control in Aquaculture, a Review of Current Knowledge and Future Perspectives. Front Microbiol 9.

18. Henriksen NNSE, Schostag MD, Balder SR, Bech PK, Strube ML, Sonnenschein EC, Gram L. 2022. The ability of *Phaeobacter inhibens* to produce tropodithietic acid influences the community dynamics of a microalgal microbiome. ISME COMMUN 2:1–11.

19. Timmerman HM, Koning CJM, Mulder L, Rombouts FM, Beynen AC. 2004. Monostrain, multistrain and multispecies probiotics—A comparison of functionality and efficacy. International Journal of Food Microbiology 96:219–233.

20. Grotkjær T, Bentzon-Tilia M, D’Alvise P, Dierckens K, Bossier P, Gram L. 2016. *Phaeobacter inhibens* as probiotic bacteria in non-axenic *Artemia* and algae cultures. Aquaculture 462:64–69.

21. Grotkjær T, Bentzon-Tilia M, D’Alvise P, Dourala N, Nielsen KF, Gram L. 2016. Isolation of TDA-producing *Phaeobacter* strains from sea bass larval rearing units and their probiotic effect against pathogenic *Vibrio* spp. in *Artemia* cultures. Systematic and Applied Microbiology 39:180–188.

22. Segev E, Wyche TP, Kim KH, Petersen J, Ellebrandt C, Vlamakis H, Barteneva N, Paulson JN, Chai L, Clardy J, Kolter R. 2016. Dynamic metabolic exchange governs a marine algal-bacterial interaction. eLife 5:e17473.

23. Biondi N, Cheloni G, Rodolfi L, Viti C, Giovannetti L, Tredici MR. 2017. *Tetraselmis suecica* F&M-M33 growth is influenced by its associated bacteria. Microb Biotechnol 11:211–223.

24. Le Reun N, Bramucci A, Ajani P, Khalil A, Raina J-B, Seymour JR. 2023. Temporal variability in the growth-enhancing effects of different bacteria within the microbiome of the diatom *Actinocyclus* sp. Front Microbiol 14.

25. Roager L, Kempen PJ, Bentzon-Tilia M, Sonnenschein EC, Gram L. 2023. Impact of host species on assembly, composition, and functional profiles of phycosphere microbiomes. bioRxiv 10.1101/2023.11.08.566273.

26. Dittmann KK, Sonnenschein EC, Egan S, Gram L, Bentzon-Tilia M. 2019. Impact of *Phaeobacter inhibens* on marine eukaryote-associated microbial communities. Environmental Microbiology Reports 11:401–413.

27. Seyedsayamdost MR, Case RJ, Kolter R, Clardy J. 2011. The Jekyll-and-Hyde chemistry of *Phaeobacter gallaeciensis*. 4. Nature Chem 3:331–335.

28. Moody SC. 2014. Microbial co-culture: harnessing intermicrobial signaling for the production of novel antimicrobials. Future Microbiology 9:575–578.

29. Scherlach K, Hertweck C. 2021. Mining and unearthing hidden biosynthetic potential. 1. Nat Commun 12:3864.

30. Selegato DM, Castro-Gamboa I. 2023. Enhancing chemical and biological diversity by co-cultivation. Frontiers in Microbiology 14.

31. Croxatto A, Lauritz J, Chen C, Milton DL. 2007. *Vibrio anguillarum* colonization of rainbow trout integument requires a DNA locus involved in exopolysaccharide transport and biosynthesis. Environ Microbiol 9:370–382.

32. Andersen JB, Sternberg C, Poulsen LK, Bjørn SP, Givskov M, Molin S. 1998. New Unstable Variants of Green Fluorescent Protein for Studies of Transient Gene Expression in Bacteria. Applied and Environmental Microbiology 64:2240–2246.

33. Guillard RRL. 1975. Culture of Phytoplankton for Feeding Marine Invertebrates, p. 29–60. *In* Smith, WL, Chanley, MH (eds.), Culture of Marine Invertebrate Animals: Proceedings — 1st Conference on Culture of Marine Invertebrate Animals Greenport. Springer US, Boston, MA.

34. Guillard RRL, Ryther JH. 1962. Studies of marine planktonic diatoms: I. *Cyclotella nana* Hustedt, and *Detonula confervacea* (cleve) Gran. Can J Microbiol 8:229–239.

35. Boström KH, Simu K, Hagström Å, Riemann L. 2004. Optimization of DNA extraction for quantitative marine bacterioplankton community analysis. Limnology and Oceanography: Methods 2:365–373.

36. Roager L, Sonnenschein EC, Gram L. 2023. Sequence-Based Characterization of Microalgal Microbiomes: Impact of DNA Extraction Protocol on Yield and Community Composition. Microbiol Spectr 11:e03408–22.

37. Bolyen E, Rideout JR, Dillon MR, Bokulich NA, Abnet CC, Al-Ghalith GA, Alexander H, Alm EJ, Arumugam M, Asnicar F, Bai Y, Bisanz JE, Bittinger K, Brejnrod A, Brislawn CJ, Brown CT, Callahan BJ, Caraballo-Rodríguez AM, Chase J, Cope EK, Da Silva R, Diener C, Dorrestein PC, Douglas GM, Durall DM, Duvallet C, Edwardson CF, Ernst M, Estaki M, Fouquier J, Gauglitz JM, Gibbons SM, Gibson DL, Gonzalez A, Gorlick K, Guo J, Hillmann B, Holmes S, Holste H, Huttenhower C, Huttley GA, Janssen S, Jarmusch AK, Jiang L, Kaehler BD, Kang KB, Keefe CR, Keim P, Kelley ST, Knights D, Koester I, Kosciolek T, Kreps J, Langille MGI, Lee J, Ley R, Liu Y-X, Loftfield E, Lozupone C, Maher M, Marotz C, Martin BD, McDonald D, McIver LJ, Melnik AV, Metcalf JL, Morgan SC, Morton JT, Naimey AT, Navas-Molina JA, Nothias LF, Orchanian SB, Pearson T, Peoples SL, Petras D, Preuss ML, Pruesse E, Rasmussen LB, Rivers A, Robeson MS, Rosenthal P, Segata N, Shaffer M, Shiffer A, Sinha R, Song SJ, Spear JR, Swafford AD, Thompson LR, Torres PJ, Trinh P, Tripathi A, Turnbaugh PJ, Ul-Hasan S, van der Hooft JJJ, Vargas F, Vázquez-Baeza Y, Vogtmann E, von Hippel M, Walters W, Wan Y, Wang M, Warren J, Weber KC, Williamson CHD, Willis AD, Xu ZZ, Zaneveld JR, Zhang Y, Zhu Q, Knight R, Caporaso JG. 2019. Reproducible, interactive, scalable and extensible microbiome data science using QIIME 2. Nat Biotechnol 37:852–857.

38. R Core Team. 2022. R: A Language and Environment for Statistical Computing. R Foundation for Statistical Computing Vienna, Austria.

39. Martin M. 2011. Cutadapt removes adapter sequences from high-throughput sequencing reads. 1. EMBnet.journal 17:10–12.

40. Callahan BJ, McMurdie PJ, Rosen MJ, Han AW, Johnson AJA, Holmes SP. 2016. DADA2: High-resolution sample inference from Illumina amplicon data. Nat Methods 13:581–583.

41. Katoh K, Standley DM. 2013. MAFFT Multiple Sequence Alignment Software Version 7: Improvements in Performance and Usability. Molecular Biology and Evolution 30:772–780.

42. Price MN, Dehal PS, Arkin AP. 2010. FastTree 2 – Approximately Maximum-Likelihood Trees for Large Alignments. PLOS ONE 5:e9490.

43. Weiss S, Xu ZZ, Peddada S, Amir A, Bittinger K, Gonzalez A, Lozupone C, Zaneveld JR, Vázquez-Baeza Y, Birmingham A, Hyde ER, Knight R. 2017. Normalization and microbial differential abundance strategies depend upon data characteristics. Microbiome 5:27.

44. Pedregosa F, Varoquaux G, Gramfort A, Michel V, Thirion B, Grisel O, Blondel M, Müller A, Nothman J, Louppe G, Prettenhofer P, Weiss R, Dubourg V, Vanderplas J, Passos A, Cournapeau D, Brucher M, Perrot M, Duchesnay É. 2018. Scikit-learn: Machine Learning in Python. arXiv:1201.0490. arXiv 10.48550/arXiv.1201.0490.

45. Bokulich NA, Kaehler BD, Rideout JR, Dillon M, Bolyen E, Knight R, Huttley GA, Gregory Caporaso J. 2018. Optimizing taxonomic classification of marker-gene amplicon sequences with QIIME 2’s q2-feature-classifier plugin. Microbiome 6:90.

46. Werner JJ, Koren O, Hugenholtz P, DeSantis TZ, Walters WA, Caporaso JG, Angenent LT, Knight R, Ley RE. 2012. Impact of training sets on classification of high-throughput bacterial 16s rRNA gene surveys. ISME J 6:94–103.

47. Bisanz JE. 2018. qiime2R: Importing QIIME2 artifacts and associated data into R sessions (v0.99). HTML.

48. RStudio Team. 2020. RStudio: Integrated Development Environment for R. RStudio, PBC Boston, MA.

49. Wickham H, Averick M, Bryan J, Chang W, McGowan LD, François R, Grolemund G, Hayes A, Henry L, Hester J, Kuhn M, Pedersen TL, Miller E, Bache SM, Müller K, Ooms J, Robinson D, Seidel DP, Spinu V, Takahashi K, Vaughan D, Wilke C, Woo K, Yutani H. 2019. Welcome to the Tidyverse. Journal of Open Source Software 4:1686.

50. Oksanen J, Blanchet FG, Kindt R, Legendre P, Minchin P, O’Hara B, Simpson G, Solymos P, Stevens H, Wagner H. 2015. Vegan: Community Ecology Package. R Package Version 22-1 2:1–2.

51. Anderson M. 2001. A new method for non-parametric multivariate analysis of variance. Austral Ecology 26:32–46.

52. Camacho C, Coulouris G, Avagyan V, Ma N, Papadopoulos J, Bealer K, Madden TL. 2009. BLAST+: architecture and applications. BMC Bioinformatics 10:421.

53. Gram L, Melchiorsen J, Bruhn JB. 2010. Antibacterial Activity of Marine Culturable Bacteria Collected from a Global Sampling of Ocean Surface Waters and Surface Swabs of Marine Organisms. Mar Biotechnol 12:439–451.

54. Rønneseth A, Castillo D, D’Alvise P, Tønnesen Ø, Haugland G, Grotkjaer T, Engell-Sørensen K, Nørremark L, Bergh Ø, Wergeland HI, Gram L. 2017. Comparative assessment of *Vibrio* virulence in marine fish larvae. J Fish Dis 40:1373–1385.

55. Petit III RA. 2024. dragonflye: Assemble bacterial isolate genomes from Nanopore reads (1.1.2). Perl.

56. Schwengers O, Jelonek L, Dieckmann MA, Beyvers S, Blom J, Goesmann A. 2021. Bakta: rapid & standardized annotation of bacterial genomes via alignment-free sequence identification (1.9.3). Microbial Genomics. Python.

57. Blin K, Shaw S, Augustijn HE, Reitz ZL, Biermann F, Alanjary M, Fetter A, Terlouw BR, Metcalf WW, Helfrich EJN, van Wezel GP, Medema MH, Weber T. 2023. antiSMASH 7.0: new and improved predictions for detection, regulation, chemical structures and visualisation. Nucleic Acids Research 51:W46–W50.

58. Alanjary M, Steinke K, Ziemert N. 2019. AutoMLST: an automated web server for generating multi-locus species trees highlighting natural product potential. Nucleic Acids Res 47:W276–W282.

59. Kokou F, Makridis P, Divanach P. 2012. Antibacterial activity in microalgae cultures. Aquaculture Research 43:1520–1527.

60. Soto-Rodriguez SA, Magallón-Servín P, López-Vela M, Nieves Soto M. 2022. Inhibitory effect of marine microalgae used in shrimp hatcheries on *Vibrio parahaemolyticus* responsible for acute hepatopancreatic necrosis disease. Aquaculture Research 53:1337–1347.

61. Austin B, Baudet E, Stobie M. 1992. Inhibition of bacterial fish pathogens by *Tetraselmis suecica*. Journal of Fish Diseases 15:55–61.

62. Ahern OM, Whittaker KA, Williams TC, Hunt DE, Rynearson TA. 2021. Host genotype structures the microbiome of a globally dispersed marine phytoplankton. Proceedings of the National Academy of Sciences 118:e2105207118.

63. Park BS, Choi W-J, Guo R, Kim H, Ki J-S. 2021. Changes in Free-Living and Particle-Associated Bacterial Communities Depending on the Growth Phases of Marine Green Algae, *Tetraselmis suecica*. 2. Journal of Marine Science and Engineering 9:171.

64. González JM, Simó R, Massana R, Covert JS, Casamayor EO, Pedrós-Alió C, Moran MA. 2000. Bacterial community structure associated with a dimethylsulfoniopropionate-producing North Atlantic algal bloom. Applied and Environmental Microbiology 66:4237–4246.

65. Steinrücken P, Jackson S, Müller O, Puntervoll P, Kleinegris D. 2023. A closer look into the microbiome of microalgal cultures. Frontiers in Microbiology 14:1108018.

66. Porsby CH, Nielsen KF, Gram L. 2008. *Phaeobacter* and *Ruegeria* Species of the *Roseobacter* Clade Colonize Separate Niches in a Danish Turbot ( *Scophthalmus maximus* )-Rearing Farm and Antagonize *Vibrio anguillarum* under Different Growth Conditions. Appl Environ Microbiol 74:7356–7364.

67. Planas M, Pérez-Lorenzo M, Hjelm M, Gram L, Uglenes Fiksdal I, Bergh Ø, Pintado J. 2006. Probiotic effect in vivo of *Roseobacter* strain 27-4 against *Vibrio anguillarum* infections in turbot (*Scophthalmus maximus* L.) larvae. Aquaculture 255:323–333.

68. Roager L, Athena-Vasileiadi D, Gram L, Sonnenschein EC. 2024. Antagonistic activity of *Phaeobacter piscinae* against the emerging fish pathogen *Vibrio crassostreae* in aquaculture feed algae. Applied and Environmental Microbiology 90:e01439–23.

69. Zhuang L, Zhang H. 2021. Utilizing cross-species co-cultures for discovery of novel natural products. Curr Opin Biotechnol 69:252–262.

70. Biswas J, Jana SK, Mandal S. 2022. Biotechnological impacts of *Halomonas*: a promising cell factory for industrially relevant biomolecules. Biotechnol Genet Eng Rev 1–30.

71. Velmurugan S, Raman K, Thanga Viji V, Donio MBS, Adlin Jenifer J, Babu MM, Citarasu T. 2013. Screening and characterization of antimicrobial secondary metabolites from *Halomonas salifodinae* MPM-TC and its *in vivo* antiviral influence on Indian white shrimp *Fenneropenaeus indicus* against WSSV challenge. Journal of King Saud University - Science 25:181–190.

72. Bibi F, Strobel GA, Naseer MI, Yasir M, Khalaf Al-Ghamdi AA, Azhar EI. 2018. Halophytes-associated endophytic and rhizospheric bacteria: diversity, antagonism and metabolite production. Biocontrol Science and Technology 28:192–213.

73. Fariq A, Yasmin A, Jamil M. 2019. Production, characterization and antimicrobial activities of bio-pigments by *Aquisalibacillus elongatus* MB592, *Salinicoccus sesuvii* MB597, and Halomonas aquamarina MB598 isolated from Khewra Salt Range, Pakistan. Extremophiles 23:435–449.

74. Lin S, Guo Y, Huang Z, Tang K, Wang X. 2023. Comparative Genomic Analysis of Cold-Water Coral-Derived *Sulfitobacter faviae*: Insights into Their Habitat Adaptation and Metabolism. 5. Marine Drugs 21:309.

75. Wang C-N, Liu Y, Wang J, Du Z-J, Wang M-Y. 2021. *Sulfitobacter algicola* sp. nov., isolated from green algae. Arch Microbiol 203:2351–2356.

76. Beiralas R, Ozer N, Segev E. 2023. Abundant *Sulfitobacter* marine bacteria protect *Emiliania huxleyi* algae from pathogenic bacteria. ISME COMMUN 3:100.

77. Pukall R, Buntefuss D, Frühling A, Rohde M, Kroppenstedt R, Burghardt J, Lebaron P, Bernard L, Stackebrandt E. 1999. *Sulfitobacter mediterraneus* sp. nov., a new sulfite-oxidizing member of the *α-Proteobacteria*. International journal of systematic bacteriology 49 Pt 2:513–9.

78. Ivanova EP, Gorshkova NM, Sawabe T, Zhukova NV, Hayashi K, Kurilenko VV, Alexeeva Y, Buljan V, Nicolau DV, Mikhailov VV, Christen R. 2004. *Sulfitobacter delicatus* sp. nov. and *Sulfitobacter dubius* sp. nov., respectively from a starfish (Stellaster equestris) and sea grass (Zostera marina). Int J Syst Evol Microbiol 54:475–480.

79. Sorokin D. 1995. *Sulfitobacter pontiacus* gen. nov., sp. nov. - A new heterotrophic bacterium from the Black Sea, specialized on sulfite oxidation. Microbiology 64:295–305.

80. Sieburth JM. 1960. Acrylic Acid, an “Antibiotic” Principle in Phaeocystis Blooms in Antarctic Waters. Science 132:676–677.

81. Raina J-B, Dinsdale EA, Willis BL, Bourne DG. 2010. Do the organic sulfur compounds DMSP and DMS drive coral microbial associations? Trends Microbiol 18:101–108.

82. Almeida JF, Marques M, Oliveira V, Egas C, Mil-Homens D, Viana R, Cleary DFR, Huang YM, Fialho AM, Teixeira MC, Gomes NCM, Costa R, Keller-Costa T. 2023. Marine sponge and octocoral-associated bacteria show versatile secondary metabolite biosynthesis potential and antimicrobial activities against human pathogens. 1. Marine Drugs 21:34.

83. Yang Q, Ge Y-M, Iqbal N, Yang X, Zhang X-L. 2021. *Sulfitobacter alexandrii* sp. nov., a new microalgae growth-promoting bacterium with exopolysaccharides bioflocculanting potential isolated from marine phycosphere. Antonie van Leeuwenhoek 114.

84. Long C, Lu X-L, Gao Y, Jiao B-H, Liu X-Y. 2011. Description of a *Sulfitobacter* Strain and Its Extracellular Cyclodipeptides. Evid Based Complement Alternat Med 2011:393752.

85. Klindworth A, Pruesse E, Schweer T, Peplies J, Quast C, Horn M, Glöckner FO. 2013. Evaluation of general 16S ribosomal RNA gene PCR primers for classical and next-generation sequencing-based diversity studies. Nucleic Acids Research 41:e1.

