## Supplementary material for "Stronger together: harnessing natural algal communities as potential probiotics for inhibition of aquaculture pathogens": Supp Tables and Figures

**S1**. List of thirty unique barcodes and primers used to amplify the V3-V4 region of the 16S rRNA gene in DNA from the microbiomes of *Isochrysis galbana* cultures. Primers and barcodes from (85).

| Barcode no. | Barcode | Forward primer (with barcode) | Reverse primer (with barcode) |
|---|---|---|---|
| 1 | TTTTAATC | TTTTAATCCCTACGGGNGGCWGCAG | TTTTAATCGACTACHVGGGTATCTAATCC |
| 2 | ATAATTAG | ATAATTAGCCTACGGGNGGCWGCAG | ATAATTAGGACTACHVGGGTATCTAATCC |
| 3 | ACCAAATT | ACCAAATTCCTACGGGNGGCWGCAG | ACCAAATTGACTACHVGGGTATCTAATCC |
| 4 | CTTATCAA | CTTATCAACCTACGGGNGGCWGCAG | CTTATCAAGACTACHVGGGTATCTAATCC |
| 5 | TGATCATT | TGATCATTCCTACGGGNGGCWGCAG | TGATCATTGACTACHVGGGTATCTAATCC |
| 6 | AGAATCTA | AGAATCTACCTACGGGNGGCWGCAG | AGAATCTAGACTACHVGGGTATCTAATCC |
| 7 | TCAAGAAA | TCAAGAAACCTACGGGNGGCWGCAG | TCAAGAAAGACTACHVGGGTATCTAATCC |
| 8 | ATCGAAAT | ATCGAAATCCTACGGGNGGCWGCAG | ATCGAAATGACTACHVGGGTATCTAATCC |
| 9 | ACATTTAC | ACATTTACCCTACGGGNGGCWGCAG | ACATTTACGACTACHVGGGTATCTAATCC |
| 10 | TAGAAAAC | TAGAAAACCCTACGGGNGGCWGCAG | TAGAAAACGACTACHVGGGTATCTAATCC |
| 11 | TTATCACC | TTATCACCCCTACGGGNGGCWGCAG | TTATCACCGACTACHVGGGTATCTAATCC |
| 12 | AATAGGGT | AATAGGGTCCTACGGGNGGCWGCAG | AATAGGGTGACTACHVGGGTATCTAATCC |
| 13 | ATTGCTGA | ATTGCTGACCTACGGGNGGCWGCAG | ATTGCTGAGACTACHVGGGTATCTAATCC |
| 14 | TGAGTTCT | TGAGTTCTCCTACGGGNGGCWGCAG | TGAGTTCTGACTACHVGGGTATCTAATCC |
| 15 | GGCTATTT | GGCTATTTCCTACGGGNGGCWGCAG | GGCTATTTGACTACHVGGGTATCTAATCC |
| 16 | CAAGAGAT | CAAGAGATCCTACGGGNGGCWGCAG | CAAGAGATGACTACHVGGGTATCTAATCC |
| 17 | GGAATACA | GGAATACACCTACGGGNGGCWGCAG | GGAATACAGACTACHVGGGTATCTAATCC |
| 18 | AAGGCAAT | AAGGCAATCCTACGGGNGGCWGCAG | AAGGCAATGACTACHVGGGTATCTAATCC |
| 19 | ACAAAACG | ACAAAACGCCTACGGGNGGCWGCAG | ACAAAACGGACTACHVGGGTATCTAATCC |
| 21 | TTGAGTGA | TTGAGTGACCTACGGGNGGCWGCAG | TTGAGTGAGACTACHVGGGTATCTAATCC |
| 22 | GCTTCTGA | GCTTCTGACCTACGGGNGGCWGCAG | GCTTCTGAGACTACHVGGGTATCTAATCC |
| 23 | GGCAAGAT | GGCAAGATCCTACGGGNGGCWGCAG | GGCAAGATGACTACHVGGGTATCTAATCC |
| 24 | GTGCTTTC | GTGCTTTCCCTACGGGNGGCWGCAG | GTGCTTTCGACTACHVGGGTATCTAATCC |
| 25 | ACACACTG | ACACACTGCCTACGGGNGGCWGCAG | ACACACTGGACTACHVGGGTATCTAATCC |
| 26 | CGATTCTG | CGATTCTGCCTACGGGNGGCWGCAG | CGATTCTGGACTACHVGGGTATCTAATCC |
| 27 | GCAGAGTT | GCAGAGTTCCTACGGGNGGCWGCAG | GCAGAGTTGACTACHVGGGTATCTAATCC |
| 30 | GCTTGGTT | GCTTGGTTCCTACGGGNGGCWGCAG | GCTTGGTTGACTACHVGGGTATCTAATCC |
| 31 | ACAGGCTT | ACAGGCTTCCTACGGGNGGCWGCAG | ACAGGCTTGACTACHVGGGTATCTAATCC |
| 40 | GAGAGGGA | GAGAGGGACCTACGGGNGGCWGCA | GAGAGGGAGACTACHVGGGTATCTAATCC |

**Table S2.** Algal and bacterial counts of different algal cultures (AXT, AXI, NT, NI and NNI) used in the *Vibrio* inhibition assay, before and after up-concentration ("Before" and "After"), and from different algal sample fractions (Full culture, FC and Filtered microbiome, FM). Counts are reported as mean ± standard deviation. In cases where standard deviation is not reported, data originated from unreplicated experimental conditions.

| Algal culture | Algal counts (log cells/mL) | | Bacterial counts (log CFU/mL) | | |
|---|---|---|---|---|---|
| | Before | After | Before | After, FC | After, FM |
| AXT | 5.2 | 7.2 | N/A | N/A | N/A |
| AXI | 5.9 | 7.2 | N/A | N/A | N/A |
| NT | 5.7 | 7.4 | 6.6 | 8.4 | 8.1 |
| NI | 6.4 ± 0.1 | 8.0 ± 0.1 | 6.0 ± 0.3 | 7.6 ± 0.2 | 7.8 ± 0.2 |
| NNI | 6.2 ± 0.6 | 7.5 ± 0.4 | 5.7 ± 0.6 | 7.7 ± 0.4 | 7.6 ± 0.3 |

**Table S3**. DNA concentration after extraction, DNA amount provided for sequencing and number of sequences obtained from each sample (after denoising and filtering). Sample names indicate replicate (“R1”, “R2”), source culture (“NI”, “NNI”), sample fraction (Full culture: “FC”, Filtered microbiome: “FM”), dilution factor of the algal microbiome (0 to -5), and starting inoculum of *V. anguillarum* NB10_gfp (“V3”, “V5” for 3 or 5 log CFU mL$^{-1}$) for samples obtained after the inhibition assay. The algal cultures were sampled before up-concentrating and filtering (“raw”, triplicates), and each fraction (FC and FM) was sampled before the enrichment assay (“bf”). Sterile media controls were included (“MB-ctrl”).

| Sample name | DNA concentration after extraction (ng/µL) | Amount of DNA provided for sequencing (ng) | Number of sequences obtained |
|---|---|---|---|
| R1-NNI-FC_0_V5 | 43.2 | 250 | 172,237 |
| R1-NNI-FC_-3_V5 | 60.8 | 250 | 659,903 |
| R1-NNI-FC_-4_V5 | 54.8 | 250 | 141,745 |
| R1-NNI-FC_-5_V3 | 36.8 | 250 | 125,582 |
| R1-NNI-FC_bf | 33.2 | 250 | 42,434 |
| R1-NNI-FM_0_V5 | 100 | 250 | 37,770 |
| R1-NNI-FM_-2_V5 | 66.4 | 250 | 283,545 |
| R1-NNI-FM_-3_V5 | 47.8 | 250 | 211,372 |
| R1-NNI-FM_-3_V3 | 77.6 | 250 | 422,921 |
| R1-NNI-FM_-4_V3 | 112 | 250 | 275,217 |
| R1-NNI-FM_bf | 4.1 | 192,7 | 175,885 |
| R1-NNI-MB-ctrl | 0.834 | 39,198 | 11,352 |
| R1-NNI-raw1 | 55.4 | 250 | 162,772 |
| R1-NNI-raw2 | 85.4 | 250 | 54,564 |
| R1-NNI-raw3 | 71.6 | 250 | 40,223 |
| R2-NNI-FC_0_V5 | 58.4 | 250 | 75,491 |
| R2-NNI-FC_-1_V5 | 39 | 250 | 550,497 |
| R2-NNI-FC_-3_V3 | 95.4 | 250 | 106,340 |
| R2-NNI-FC_-5_V3 | 73.4 | 250 | 324,687 |
| R2-NNI-FC_bf | 85 | 250 | 62,636 |
| R2-NNI-FM_0_V5 | 75.4 | 250 | 238,018 |
| R2-NNI-FM_-3_V3 | 120 | 250 | 162,818 |

| | | | |
|---|---|---|---|
| R2-NNI-FM_-4_V3 | 48 | 250 | 294,405 |
| R2-NNI-FM_bf | 17.3 | 250 | 145,227 |
| R2-NNI-raw1 | 43.4 | 250 | 25,781 |
| R2-NNI-raw2 | 72.4 | 250 | 49,342 |
| R2-NNI-raw3 | 72.8 | 250 | 74,725 |
| R2-NI-FC_0_V5 | 81.2 | 250 | 291,271 |
| R2-NI-FC_-2_V5 | 16.2 | 250 | 391,227 |
| R2-NI-FC_-3_V3 | 61.6 | 250 | 297,668 |
| R2-NI-FC_-4_V3 | 74.4 | 250 | 224,317 |
| R2-NI-FC_bf | 120 | 250 | 33,188 |
| R2-NI-FM_0_V5 | 95.2 | 250 | 474,932 |
| R2-NI-FM_-1_V5 | 61 | 250 | 265,940 |
| R2-NI-FM_-2_V3 | 75 | 250 | 229,292 |
| R2-NI-FM_-4_V3 | 50 | 250 | 226,214 |
| R2-NI-FM_bf | 16.4 | 250 | 367,349 |
| R2-NI-MB-ctrl | 1.98 | 93,06 | 39,985 |
| R2-NI-raw1 | 86.8 | 250 | 43,457 |
| R2-NI-raw2 | 92.2 | 250 | 169,394 |
| R2-NI-raw3 | 73.4 | 250 | 409,195 |

**Table S4.** 64 bacterial strains isolated from the native („Raw culture", 19 strains) or enriched inhibitory („Full culture": FC, or „Filtered microbiome": FM, 45 strains) microbiome of *Isochrysis galbana*, and tentatively identified using full-length 16S rRNA gene sequencing. The isolates originate from two different *I. galbana* cultures (NI and NNI), acquired at different times from the same aquaculture unit.

| Isolate | Source | Best BLAST hit | Query Cover | E value | ID% |
|---|---|---|---|---|---|
| A1 | Raw culture, NNI | *Alteromonas marina* | 99% | 0.0 | 98.14% |
| A2 | Raw culture, NNI | *Sulfitobacter pacificus* | 99% | 0.0 | 99.17% |
| A3 | Raw culture, NNI | *Roseovarius nubinhibens* | 99% | 0.0 | 98.61% |
| A4 | Raw culture, NNI | *Roseovarius nubinhibens* | 100% | 0.0 | 98.88% |
| A5 | Raw culture, NNI | *Sulfitobacter* sp. | 100% | 0.0 | 98.63% |
| A6 | Raw culture, NNI | *Sulfitobacter pacificus* | 100% | 0.0 | 99.71% |
| A7 | Raw culture, NNI | *Qipengyuania nanhaisediminis* | 100% | 0.0 | 99.54% |
| A8 | Raw culture, NNI | *Qipengyuania aquimaris* | 99% | 0.0 | 98.90% |
| A9 | Raw culture, NNI | *Ruegeria* sp. | 100% | 0.0 | 98.29% |
| B1 | Raw culture, NNI | *Alteromonas macleodii* | 100% | 0.0 | 97.96% |
| B2 | Raw culture, NNI | *Qipengyuania xiamenensis* | 99% | 0.0 | 99.56% |
| B3 | Raw culture, NI | *Croceibacter* sp. | 100% | 0.0 | 99.22% |
| B4 | Raw culture, NI | *Mameliella alba* | 99% | 0.0 | 98.43% |
| B5 | Raw culture, NI | *Sulfitobacter* sp. LZD014 | 99% | 0.0 | 99.34% |
| B6 | Raw culture, NI | *Sulfitobacter* sp. | 100% | 0.0 | 98.82% |
| B7 | Raw culture, NI | *Phaeobacter piscinae* | 100% | 0.0 | 99.57% |
| B9 | Raw culture, NI | *Phaeobacter piscinae* | 99% | 0.0 | 99.43% |
| C1 | Raw culture, NI | *Sulfitobacter* sp. | 99% | 0.0 | 99.57% |
| C2 | FC, NNI | *Roseovarius nubinhibens* | 99% | 0.0 | 99.31% |
| C3 | FC, NNI | *Phaeobacter piscinae* | 99% | 0.0 | 99.43% |
| C5 | FC, NNI | *Roseovarius nubinhibens* | 100% | 0.0 | 99.37% |
| C6 | FC, NNI | *Sulfitobacter* sp. LZD014 | 100% | 0.0 | 98.86% |
| C7 | FC, NNI | *Roseovarius nubinhibens* | 100% | 0.0 | 99.62% |
| C8 | FC, NNI | *Alteromonas tagae* | 99% | 0.0 | 97.35% |
| C9 | FC, NNI | *Sulfitobacter pacificus* | 99% | 0.0 | 99.24% |
| D1 | FC, NNI | *Sulfitobacter* sp. LZD014 | 99% | 0.0 | 99.52% |
| D2 | FC, NNI | *Halomonas alkaliphila* | 99% | 0.0 | 98.89% |

| | | | | | |
|---|---|---|---|---|---|
| D3 | FC, NNI | *Sulfitobacter* sp. | 99% | 0.0 | 97.53% |
| D4 | FC, NNI | *Phaeobacter piscinae* | 99% | 0.0 | 99.14% |
| D5 | FC, NNI | *Alteromonas* sp. | 98% | 0.0 | 98.10% |
| D6 | FC, NI | *Phaeobacter piscinae* | 100% | 0.0 | 99.24% |
| E2 | FC, NI | *Alteromonas macleodii* | 99% | 0.0 | 99.14% |
| E3 | FC, NI | *Phaeobacter piscinae* | 99% | 0.0 | 99.05% |
| E4 | FC, NI | *Phaeobacter piscinae* | 100% | 0.0 | 99.42% |
| E5 | FC, NI | *Sulfitobacter* sp. LZD014 | 100% | 0.0 | 99.33% |
| E6 | FC, NI | *Phaeobacter* sp. M8-4.3 | 99% | 0.0 | 98.24% |
| E7 | FC, NI | *Sulfitobacter* sp. LZD014 | 99% | 0.0 | 99.23% |
| E8 | FC, NI | *Phaeobacter piscinae* | 100% | 0.0 | 99.42% |
| E9 | FC, NI | *Phaeobacter piscinae* | 100% | 0.0 | 98.90% |
| F1 | FC, NI | *Sulfitobacter* sp. | 100% | 0.0 | 98.82% |
| F2 | FC, NI | *Vibrio anguillarum* | 100% | 0.0 | 97.43% |
| F3 | FC, NI | *Phaeobacter* sp. M8-4.3 | 99% | 0.0 | 98.07% |
| F4 | FC, NI | *Sulfitobacter* sp. | 94% | 0.0 | 96.22% |
| F5 | FM, NNI | *Vibrio* sp. | 100% | 0.0 | 98.70% |
| F6 | FM, NNI | *Phaeobacter piscinae* | 100% | 0.0 | 98.72% |
| F7 | FM, NNI | *Sulfitobacter* sp. | 99% | 0.0 | 99.33% |
| F8 | FM, NNI | *Roseovarius nubinhibens* | 99% | 0.0 | 99.44% |
| F9 | FM, NNI | *Roseovarius nubinhibens* | 99% | 0.0 | 99.71% |
| G1 | FM, NNI | *Sulfitobacter* sp. | 99% | 0.0 | 99.43% |
| G2 | FM, NNI | *Phaeobacter piscinae* | 100% | 0.0 | 99.23% |
| G3 | FM, NNI | *Alteromonas* sp. | 97% | 0.0 | 97.75% |
| G5 | FM, NNI | *Roseovarius nubinhibens* | 100% | 0.0 | 98.91% |
| G7 | FM, NNI | *Vibrio* sp. | 99% | 0.0 | 97.46% |
| G8 | FM, NNI | *Roseovarius nubinhibens* | 99% | 0.0 | 99.05% |
| H1 | FM, NI | *Sulfitobacter* sp. LZD014 | 99% | 0.0 | 99.61% |
| H2 | FM, NI | *Phaeobacter piscinae* | 100% | 0.0 | 98.96% |
| H3 | FM, NI | *Phaeobacter piscinae* | 99% | 0.0 | 99.33% |
| H4 | FM, NI | *Phaeobacter* sp. M8-4.3 | 99% | 0.0 | 98.33% |
| H8 | FM, NI | *Phaeobacter piscinae* | 99% | 0.0 | 99.33% |
| H9 | FM, NI | *Alteromonas macleodii* | 100% | 0.0 | 91.23% |
| I1 | FM, NI | *Phaeobacter piscinae* | 100% | 0.0 | 99.14% |

| | | | | | |
|---|---|---|---|---|---|
| I2 | FM, NI | *Phaeobacter piscinae* | 99% | 0.0 | 99.04% |
| I3 | FM, NI | *Phaeobacter piscinae* | 100% | 0.0 | 99.24% |
| I4 | Raw culture, NI | *Phaeobacter piscinae* | 99% | 0.0 | 98.94% |

**Table S5.** Inhibition of nine *V. anguillarum* strains by the co-culture of *Halomonas campaniensis* (isolate D2) and *Sulfitobacter pontiacus* (isolate D3). *Phaeobacter piscinae* isolate H2 was used as positive control. Clearing zone of different degrees (+, ++, +++), a faint clearing zone (*), or no clearing zone (-) after 24 hours of incubation. Virulence ranks as defined by Rønneseth et al (54).

| ***V. anguillarum* strain** | **Virulence rank** | **Inhibition by *H. campaniensis* D2 monoculture** | **Inhibition by *S. pontiacus* D3 monoculture** | **Inhibition by co-culture of *H. campaniensis* D2 and *S. pontiacus* D3** | **Inhibition by *P. piscinae* H2** |
|---|---|---|---|---|---|
| 90-11-286 | High | - | - | - | +++ |
| DSM21597 | High | - | - | - | +++ |
| PF7 | High | - | - | * | +++ |
| PF4 | High | - | - | - | +++ |
| 9014/8 | Medium | - | - | - | +++ |
| S2 2/9 | Medium | - | - | - | +++ |
| 4299 | Low | * | - | ++ | +++ |
| NB10 | Low | * | - | ++ | +++ |
| 775 | N/A | + | - | + | +++ |

**Table S6.** Biosynthetic gene clusters (BGCs) predicted by antiSMASH 7.0 in the genomes of *Halomonas campaniensis* D2 and *Sulfitobacter pontiacus* D3.

| Strain | Cluster | Type of BGC | Most similar known BGC in MIBiG | Similarity |
|---|---|---|---|---|
| *H. campaniensis* D2 | 1 | NI-siderophore | BGC0001572 | 66% |
| | 2 | ranthipeptide | BGC0000413 | 2% |
| | 3 | redox-cofactor | BGC0001131 | 25% |
| | 4 | RiPP-like | / | / |
| | 5 | T1PKS | / | / |
| | 6 | betalactone | BGC0001103 | 20% |
| | 7 | ectoine | BGC0000859 | 75% |
| *S. pontiacus* D3 | 1 | RiPP-like | / | / |
| | 2 | betalactone | / | / |
| | 3 | hserlactone | / | / |
| | 4 | redox-cofactor | / | / |

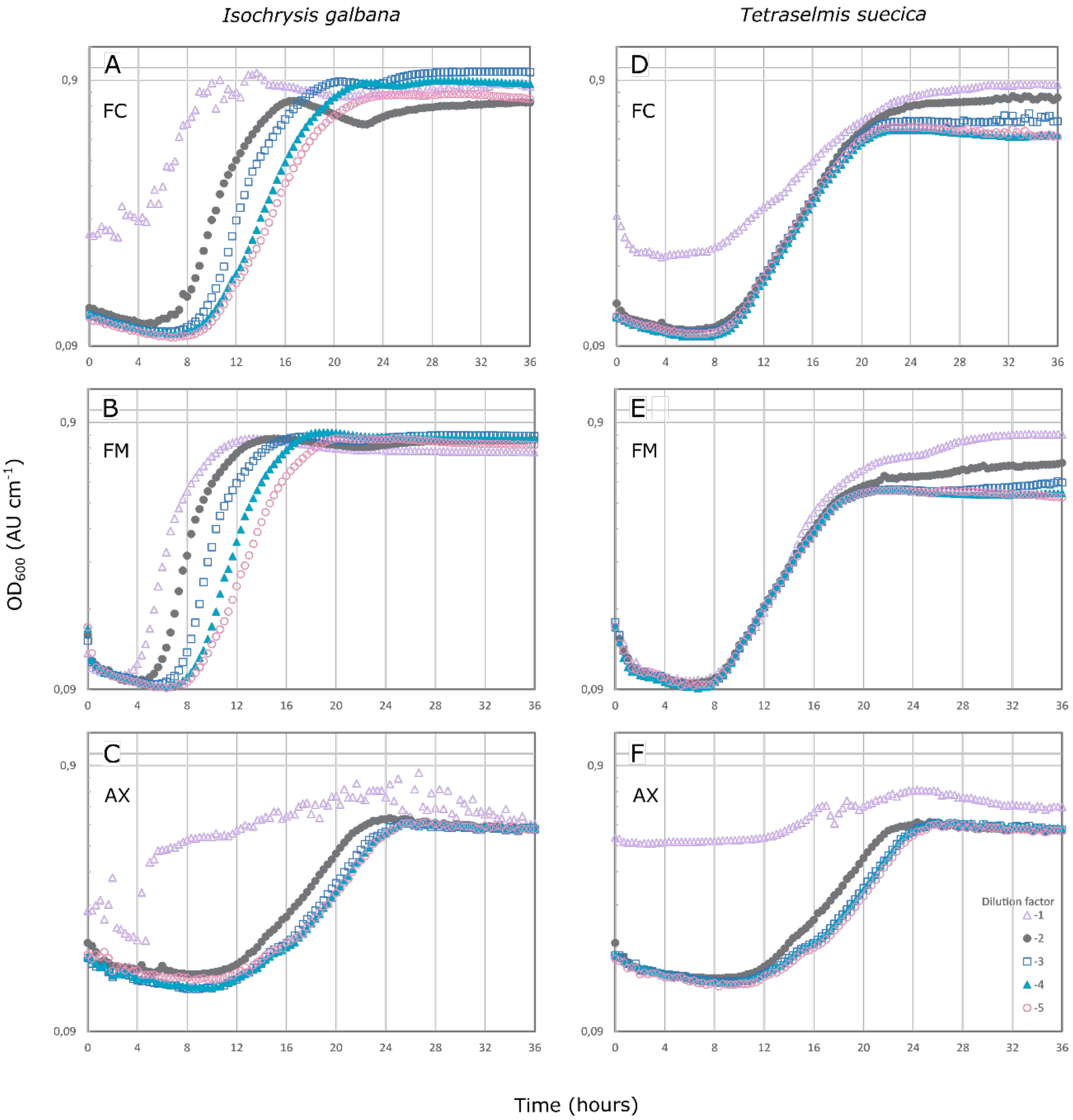


**Figure S1.** Inhibition assay by *Isochrysis galbana* (left) and *Tetraselmis suecica* (right) microbiomes against *Vibrio anguillarum* NB10_gfp, at a starting concentration of 3.1 ± 0.3 log CFU $mL^{-1}$, as measured by absorbance at 600 nm. The inhibitory effect of serial dilutions ($10^{-1}$ (△), $10^{-2}$ (●), $10^{-3}$ (□), $10^{-4}$ (▲), $10^{-5}$ (○)) of different fractions of the algal cultures have been tested: full culture (FC; S1A and S1D), with algal and bacterial cells; filtered microbiome (FM; S1B and S1E), where algal cells have been removed; and axenic (AX; S1C and S1F) cultures, where the algae cells are free of bacteria.

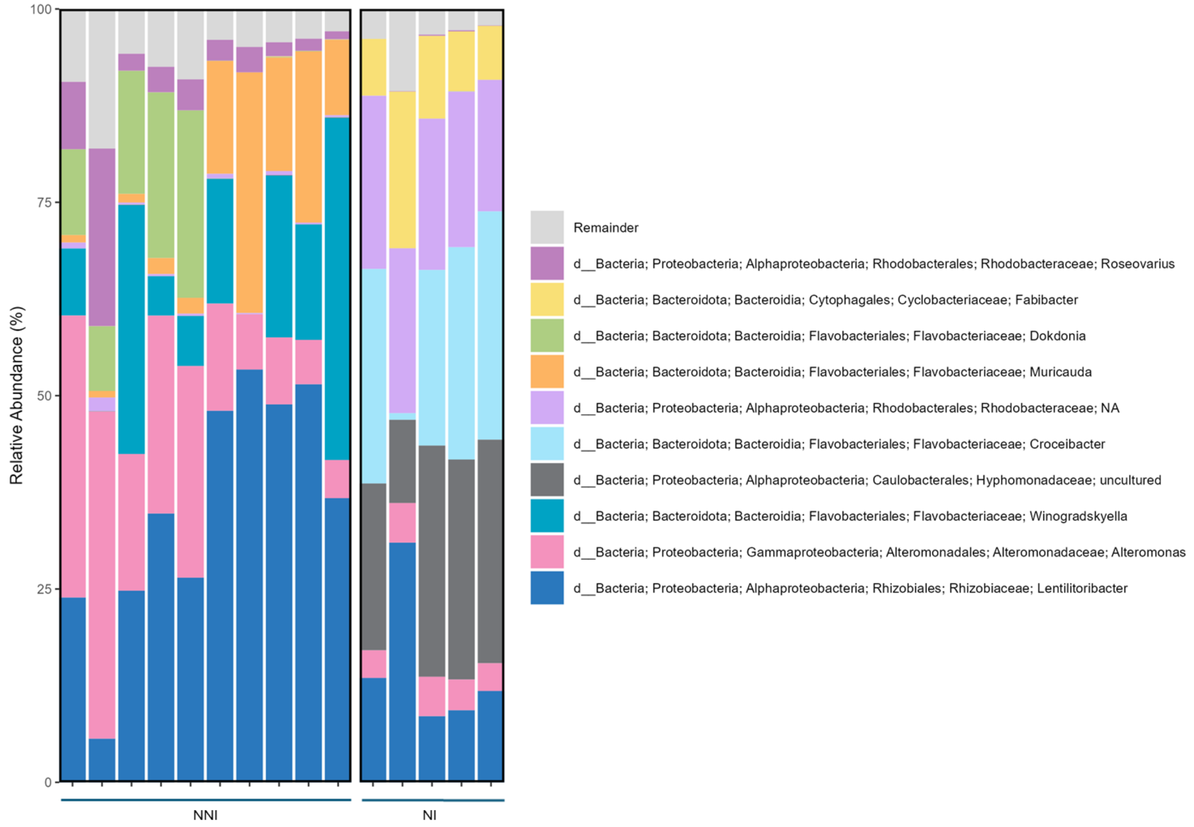


**Figure S2.** Native microbiome composition (top 10 most abundant genera) of two *Isochrysis galbana* cultures of different age based on the 16S rRNA amplicon sequencing results. One culture, NNI, has been newly provided by an aquaculture facility and has a relatively freshly recruited microbiome (left). The other culture, NI has been regularly subcultured under laboratory conditions for almost four years (right).

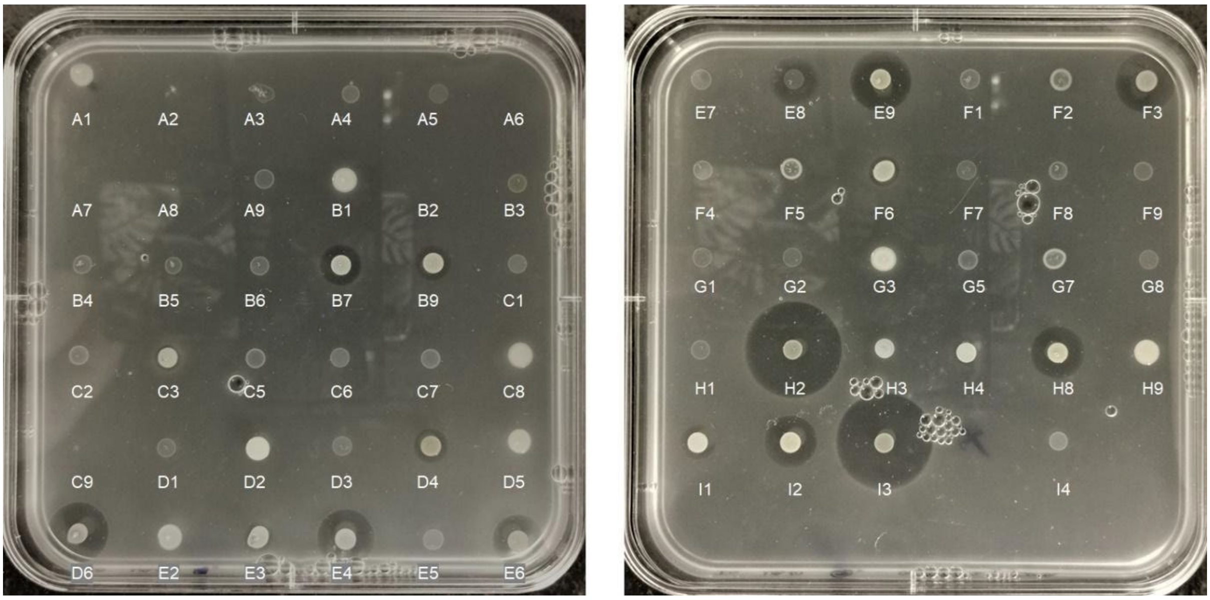


**Figure S3.** Inhibition of *Vibrio anguillarum* NB10_gfp by 64 isolates from the *Isochrysis galbana* microbiome.
